# Machine Learning and Optimal Control of Enzyme Activities to Preserve Solvent Capacity in the Cell

**DOI:** 10.1101/2020.04.06.028035

**Authors:** Samuel Britton, Mark Alber, William R. Cannon

## Abstract

Experimental measurements or computational model predictions of the post-translational regulation of enzymes needed in a metabolic pathway is a difficult problem. Consequently, regulation is mostly known only for well-studied reactions of central metabolism in various model organisms. In this study, we utilize two approaches to predict enzyme regulation policies and investigate the hypothesis that regulation is driven by the need to maintain the solvent capacity in the cell. The first predictive method uses a statistical thermodynamics and metabolic control theory framework while the second method is performed using a hybrid optimization-reinforcement learning approach. Efficient regulation schemes were learned from experimental data that either agree with theoretical calculations or result in a higher cell fitness using maximum useful work as a metric. Model predictions provide the following novel general principles: (1) the regulation itself causes the reactions to be much further from equilibrium instead of the common assumption that highly non-equilibrium reactions are the targets for regulation; (2) regulation is used to maintain the concentrations of both immediate and downstream product concentrations rather than to maintain a specific energy charge; and (3) the minimal regulation needed to maintain metabolite levels at physiological concentrations also results in the maximal energy production rate that can be obtained at physiological conditions. The resulting energy production rate is an emergent property of regulation which may be represented by a high value of the adenylate energy charge. In addition, the predictions demonstrate that the amount of regulation needed can be minimized if it is applied at the beginning or branch point of a pathway, in agreement with common notions. The approach is demonstrated for three pathways in the central metabolism of *E. coli* (gluconeogenesis, glycolysis-TCA and Pentose Phosphate-TCA) that each require different regulation schemes. It is shown quantitatively that hexokinase, glucose 6-phosphate dehydrogenase and glyceraldehyde phosphate dehydrogenase, all branch points of pathways, play the largest roles in regulating central metabolism.

## Introduction

While our understanding of regulation of transcription and post-transcriptional processes has blossomed in the past 25 years due to advances in high-throughput experimental technologies such as RNA expression, ChIP-Seq, and mass spectrometry-based proteomics, our understanding of post-translational regulation has advanced^1–4^, but not as rapidly or as far.

Recent breakthroughs include work in which mass spectrometry and NMR measured metabolite and protein levels, along with fluxes modeled from 13C isotope labeling were used with Michaelis-Menten kinetics to determine whether the predicted reaction fluxes matched fluxes modeled from isotope labeling data^2^. The correlation between predicted fluxes were evaluated with and without regulation. If the match was better with regulation, then regulation was assumed. The work was a *tour de force* in that chemostat studies we used to carefully measure both absolute and relative metabolomics data while at the same time cover as much of the proteome as possible. In addition, Michaelis-Menten kinetic models addressed multiple levels of regulation. The payoff was not only predictions of which enzymes might be regulated, but also inferences about the regulating molecule.

In addressing possible scalability (or at least cost of experimentation) in the previously mentioned study, a similarly sophisticated informatics approach was used to develop a model of small molecule regulatory networks from curated databases of enzymes, integrate the regulatory network with a metabolic model of *E. coli*, and distill information on how substrates and inhibitors contribute to metabolic flux regulation^3^. Interestingly, this work did not find support for the common notion that reactions which are furthest from equilibrium are those that are most likely regulated.

Fifty years ago it was postulated that the purpose of post-translational regulation in metabolism is to either maintain a balance of the energy charge of the adenylate pool^5^, or to control solvent properties^6^. Solvent properties have long been recognized as important determinants of cellular activity and function. Atkinson recognized that the maintenance of physiological concentrations of metabolites may well be the most pressing problem of metabolic control^6^. Metabolite concentrations are exponential functions of the standard chemical potentials but only a linear function of the rate constants. Consequently, metabolite concentrations are less a function of the reaction kinetics and primarily a function of a molecule’s standard chemical potential, which varies over a small range across species because solution conditions inside a cell also vary over a small range. Interestingly, the set of enzymes which are post-translationally regulated is relatively well-conserved across species as well^3^, despite the fact that the rate constants for the same enzymes can vary dramatically^7^.

In addition to metabolite concentrations *per se,* solvent capacity in the cell has recently focused on molecular crowding^8,9^ and the impairment of diffusion^10^. As a cell approaches equilibrium, the cell’s cytoplasm can become glassy such that diffusion is limited. At the same time, control of metabolites through regulation of enzyme activities is no longer effective near equilibrium^11^. The equilibrium constant *K* for a reaction is the ratio of the exponent of the standard chemical potentials. Consequently, metabolite concentrations may potentially approach values determined by their standard chemical potentials in solution, which can be quite large for highly charged metabolites like fructose 1,6-bisphosphate and acetyl-coenzyme A. Not only will metabolite levels rise, but also less water will be produced by metabolism inside the cell. In *E. coli,* up to 50% of the bulk water is produced by metabolism^12^. Even away from equilibrium, cells clearly must regulate metabolite levels to prevent high concentrations that would be detrimental to diffusional processes necessary for life.

Here, we investigate the hypothesis that the post-translational regulation of enzymes is at least in part driven by the need to maintain the solvent capacity in the cell. We evaluate this hypothesis by comparing experimental metabolomics data with steady state concentrations predicted computationally from equations for reformulated mass action kinetics. Using quantitative metabolomics data as well as physical and biological principles, metabolic control analysis and alternatively reinforcement learning are used to predict the control of activity required to bring metabolite levels down to observed values. Consequently, the machine learning results confirm that an optimal control policy can be formulated which directly achieves minimal regulation by efficiently reducing excessive metabolite concentrations.

The predictions agree with known regulation of central metabolism in model organisms. Moreover, these results show that regulated enzymes have higher free energies of reaction precisely because of the regulation, turning common wisdom about enzyme regulation upside-down. Instead of highly non-equilibrium reactions being the targets for regulation in metabolic pathways^13,14^, regulation results in reactions being much further from equilibrium than non-regulated reactions. Being further away from equilibrium than other reactions is an effect, not a cause, of regulation.

## Results

We solve the prediction problem of which enzyme to regulate by a novel combination of methods from statistical thermodynamics, control theory and reinforcement learning (RL). The initial step is to determine steady state concentrations without applying regulation by using numerical optimization of the respective ordinary differential equations on a convex energy surface. The convex energy surface for metabolic dynamics is obtained by assuming that the time dependence is the same for all reactions in the Marcelin-de Donder dynamical force equation for mass action kinetics^15^. Due to the assumption that the reactions all occur on the same time scale, the thermodynamic odds of each reaction (Methods, Eqn. (7)) are similar in value in upper glycolysis, lower glycolysis and the TCA cycle, though varying by a factor of two due to stoichiometry. Such a configuration is known as a maximum reaction path entropy configuration^16,17^. Figure 2 shows the resulting steady state reaction fluxes and reaction free energies for the glycolysis-PPP-TCA cycle under high NAD/NADH and low NADP/NADPH conditions.

If there are no constraints, the maximum path entropy configuration also results in a maximal entropy distribution of metabolites. However, the metabolites will be constrained to be away from the equilibrium distribution if there are nonequilibrium boundary conditions. Since the initially predicted concentrations will then be proportional to their Boltzmann probabilities, the initially predicted concentrations may be exceedingly high^6^ compared to experimentally observed values from isotope-labeled, mass spectrometry measurements^18,19^. However, these high concentrations allow for highly effective inference of regulation to control the concentrations. The predicted concentrations, *ñ_i_*, are brought into alignment with experimental observations, *n_i_*, by applying regulation. Regulation is determined using either a Metabolic Control Analysis (MCA) approach, or a hybrid optimization-reinforcement learning (RL) approach (Methods). In both cases, regulation is applied in the form of an activity coefficient, *α_j_*, that scales the reaction flux for reaction *j*, where *α_j_*, = 1.0 indicates no regulation while *α_j_*, = 0.0 indicates complete regulation.

In the two MCA based methods that were developed, reactions are regulated based on the sensitivity of the predicted concentrations to the activity coefficient that modulates each reaction, which is carried out by a specific enzyme. The sensitivity of the *i^th^* metabolite with concentration *n_i_* (observed or predicted) to the activity, *α_j_*, of enzyme *j*, is described by the concentration control coefficient, 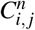,

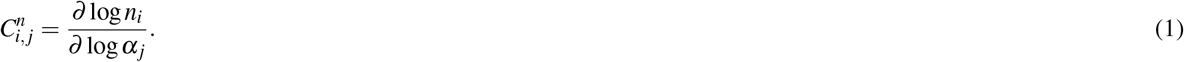

When using predicted concentrations, *ñ_i_*, we write 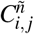 to specify the concentration control coefficient for predicted metabolite concentrations. We utilize a loss function defined as the logarithm of the division of the predicted concentrations or counts to the measured concentrations or counts, *L_i_*, = log(*ñ_i_*,/*n_i_*). The change in the loss function due to a change in the activity of reaction *j* is

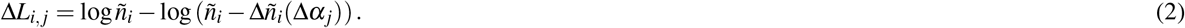

The reaction *j* selected for regulation is the one whose change in activity results in the largest change in the loss functions of all metabolites whose predicted concentrations exceed the experimentally observed concentrations, as determined by Δ*L_j_* = Σ_*i*_Δ*L_i,j_*. Regulation is considered complete when predicted metabolite concentrations are brought into agreement with experimental measurements.

Two approaches were taken with MCA: an unrestricted control approach in which any enzyme could be a regulator for any metabolite, and a restricted approach in which only enzymes whose immediate products exceeded the target values could be considered as a regulator. While the unrestricted approach optimizes the system as a whole, the restricted approach is consistent with the concept of modularity in biological systems. We refer to the latter as a local-control approach (MCA Local) since an enzyme’s immediate products (and possibly other metabolites) are being controlled.

The RL method (Figure 1) formulates the problem of regulation in terms of a Markov decision process^20^, which is commonly represented as a tuple {*S, A, P, R*}, where *S* represents the set of possible states (enzyme activities for each reaction), *A* represents the set of possible actions (reactions to regulate), *P* represents the transitional probabilities between states, and R represents the reward function. Reinforcement learning is utilized to obtain an optimal regulation scheme by learning from delayed environmental feedback^21,22^. Figure 1 illustrates how reactions are chosen using a policy function which returns the reaction to be regulated (action) given the current enzyme activities (state). Learning is performed by iteratively updating the state value function using environmental feedback (rewards) from solving the optimization routine. Specifically, we utilize a temporal difference bootstrapping technique^23^ called n-step SARSA^24,25^, an on-policy version of the recently popularized Q-learning method^26^. In the Methods and Supplementary Material, we provide in-depth descriptions of the theories and approaches behind the steady state optimization, the MCA methods and the RL method.

**Figure 1.**
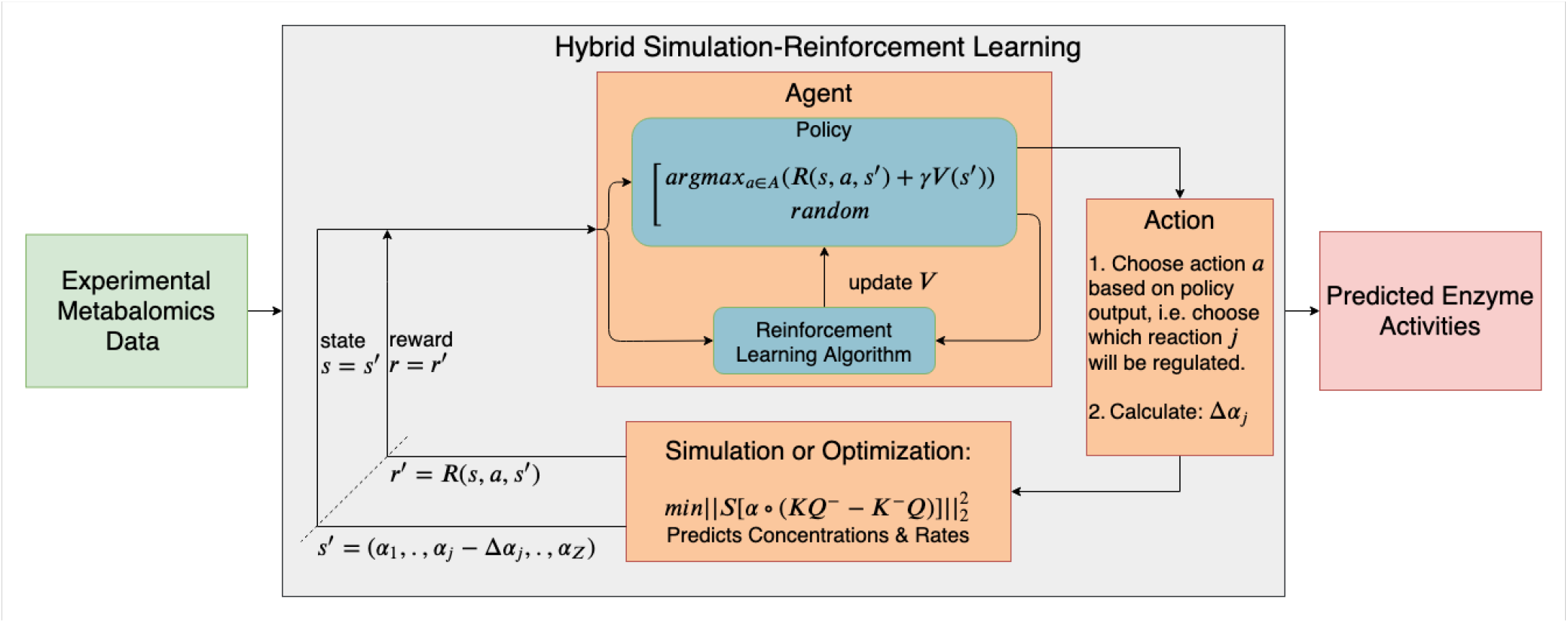
Schematic of *in silico* framework for learning regulation (grey box) with coupled simulation or optimization routine controlling environmental feedback. Initial framework input (green box) consists of target metabolite concentrations from experimental data. The output (red box) consists of a learned optimal enzyme regulation scheme necessary to reach the target concentrations. Learning is performed by repeatedly testing different regulation schemes and updating the value function, *V*, that returns a scalar value for a given set of enzyme activities. Enzyme activities, represented as states, are chosen for regulation by performing actions that are determined by a policy function. A given policy is determined by *V*. The new steady state metabolite concentrations resulting from applied regulation are determined by an optimization routine. Alterations in metabolite concentrations are a direct result of moving into a state *s*’ from a state s after taking action *a*, i.e. performing regulation. These dynamic changes are used to define a reward function, *R*, that determines environmental feedback. Rewards are used to direct the agent as it explores and learns a policy that predicts optimal enzyme regulation.

We compare the three different regulation approaches by statistically characterizing the rate of energy flow across the reactions. The rate that energy is produced in metabolism has long been known to be one of the most significant factors in metabolic regulation^5^. The sum of the rate of free energy generated across all reactions is the free energy dissipation rate, or equivalently the negative of the entropy production rate. The free energy of the *j^th^* reaction at steady state is 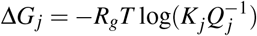, where *R_g_* is the gas constant, and *T* is the temperature, *K_j_* is the equilibrium constant and *Q_j_* is the reaction quotient. The free energy dissipation rate is defined as the rate at which free energy is dissipated^27,28^,

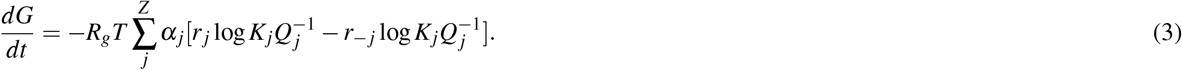

In the maximum path entropy formulation (Methods, Eqn. (10)), the rate *r_j_* is proportional to the thermodynamic driving force on the reaction, 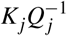. The free energy change for a reaction *j* can be broken down into two components, an energy change, Δ*E_j_* = −*R_g_T*log*K_j_*, and a configurational entropy change, *T*Δ*S_j_* = *R_g_T*log*Q_j_*^17^. As the reactions occur, the system moves towards equilibrium, decreasing the reactants and increasing the products, which results in a change in the configurational entropy due to changes in the reaction quotients. In a steady state or pseudo-steady state system, the steady state is replenished by additional nutrients such that the reaction quotients, *Q_j_*, return to their steady state values. Replenishing the steady state, however, requires work. Since the net entropy change in a pseudo-steady state system must be zero, the measure of work available for processes other than maintaining the steady state, such as replication, is,

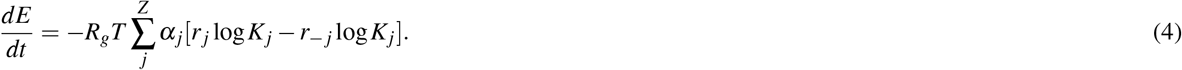

Both *dG/dt* and *dE/dt* (the energy dissipation rate) are important metrics of the rate of work produced by metabolism. When regulating reactions, a biological system must find a balance between a free energy dissipation rate that extracts energy from the environment as quickly as possible and a low rate of entropy change to maintain the pseudo-steady state. In principle, any individual or species in a pseudo-steady state that maximizes the rate of usable work, *dE*/*dt*, will outcompete those with lower rates of net work and will be the organism selected by nature.

We evaluated three different versions of *E. coli* central metabolism under four different nutrient conditions. The three different versions of metabolism were (1) gluconeogenesis, (2) glycolysis and the TCA cycle, and (3) glycolysis, the pentose phosphate pathway (PPP) and the TCA cycle (glycolysis-PPP-TCA). Metabolite concentration data used in the analysis were from *E. coli* in exponential growth with glucose as the carbon source^18,19^. In all cases, the predicted regulation matched known regulation points in central metabolism or were adjacent to known regulation points.

Below, we discuss the largest network, glycolysis-PPP-TCA, under two identical nutrient conditions except for the NADP/NADPH ratio, which is held fixed but at different values throughout each analysis. In condition 1, the NAD/NADH ratio is high (31.3) and the NADP/NADPH ratio is low (0.02), which favors flux through upper glycolysis rather than PPP. In condition 2, the NADP/NADPH ratio is also high such that NADP/NADPH = NAD/NADH = 31.3^18^. The latter condition favors increased flux through PPP. Analyses of gluconeogenesis and glycolysis and the TCA cycle are included in the supplementary materials (Figure S2 and S3). In all conditions, we compare regulation that is found by the reinforcement learning method with that found by deterministic methods using only MCA.

### High NAD/NADH require regulation of metabolite levels in glycolysis

Prediction of enzyme activities using MCA methods are deterministic. Given the conditions for fixed metabolites in which the NAD/NADP ratio is high and the NADP/NADPH ratio is low, flux is favored through upper glycolysis over PPP, and the local MCA method predicts (Figure 3A, red ‘plus’) that five reactions in glycolysis are regulated due to the enzymes hexose kinase (HEX1), phosphofructokinase (PFK), glyceraldehyde-3-phosphate dehydrogenase (GAPD), phosphoglycerate kinase (PGK), and pyruvate dehydrogenase (PDH), while one enzyme in PPP is regulated, phosphogluconolactonase (PGL), near the beginning of the pathway. It is known that regulation of PPP occurs one enzyme up from PGL at glucose 6-phosphate dehydrogenase (G6PDH) instead. But the metabolite that is over produced and is predicted to have high concentration without regulation is phosphogluconate, the product of the PGL reaction. In practice, PGL may be a hard reaction to allosterically regulate since it is a unimolecular ring opening reaction that may be catalyzed significantly by binding alone^29^.

The RL and unrestricted MCA methods both predict the same minimal regulation at HEX1 and GAPD to achieve the same goal of maintaining the predicted concentrations at or below the experimentally observed values. The RL method, however, additionally regulates PGK, pyruvate kinase (PYK), the pyruvate mitochondrial transporter (PYRt2m) and PDH to obtain a similar energy dissipation rate. As shown in Figure 3A, four of these enzymes were also regulated in the local MCA method. The difference is that HEX1 and GAPD are more extensively regulated in the RL and the unrestricted MCA methods. Despite these differences in regulation, each regulated enzyme with the exception of the pyruvate transporter are known sites of regulation (known sites of regulation are highlighted in bold). Regulation of the pyruvate transporter was only predicted in the stochastic RL approach. It is likely that this regulation should be assigned to PYK or PDH as it was in the deterministic MCA approach.

As shown in Figure 3B, whenever regulation is applied in the form of reducing the activity coefficient, the free energy of the reaction becomes more favorable compared to reactions in the same pathway (e.g., compare to the consistency of free energy changes in upper glycolysis, PPP, lower glycolysis and TCA cycle in Fig 2). Reducing the activity of an enzyme in a non-equilibrium setting will cause the reactants to increase in concentration and the products to decrease in concentration, resulting in reaction free energies being further away from equilibrium. Despite the different sites of regulation and the difference in reaction free energies for the three methods, the free energy and energy dissipation rates are similar and are the most favorable rates found (Figure 3C).

**Figure 2.**
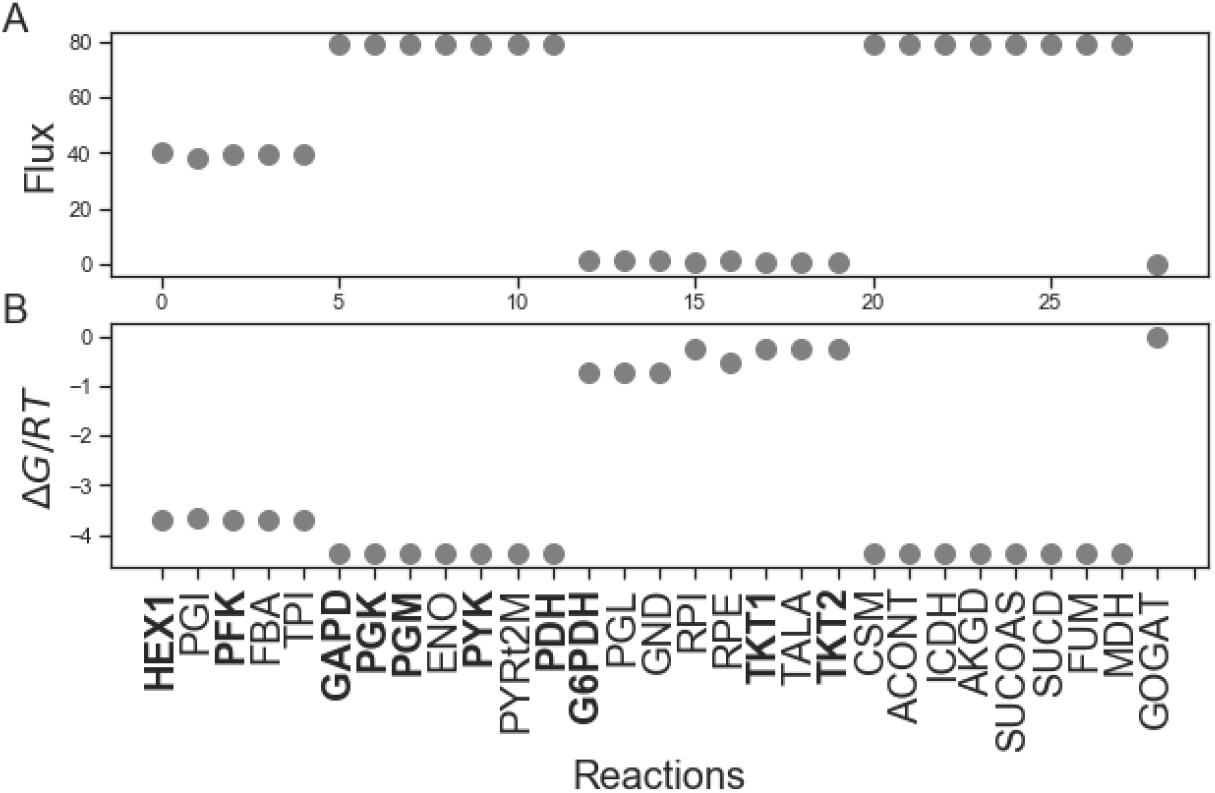
Initial steady state properties before any regulation is applied in the form of reduced activity coefficients for glycolysis-PPP-TCA cycle with high NAD/NADH and low NADP/NADPH conditions. The steady state is determined by maximizing the reaction path entropy such that the net thermodynamic driving force on each reaction is proportioned according to the governing equation for metabolite kinetics, Eqn. (14). (A) Unregulated reaction fluxes. (B) Unregulated reaction free energies. Reduction of activity coefficients to values less than 1.0 reduces both the steady state fluxes and the reaction free energies (Fig. 3–5).

### High NAD/NADH & High NADP/NADPH require additional regulation in PPP

In the second set of conditions, the NADP/NADPH ratio is also high, which in principle favors more flux through PPP. The resulting regulation is similar to the first conditions in which NADP/NADPH is low with a few exceptions (Figure 4A and 4B). The local MCA method additionally regulated G6PDH, the entry point into the PPP as well as transketolase (TKT), while the RL method no longer regulated PYK and regulated the pyruvate mitochondrial transporter (PYRt2m) rather than PDH. The latter is likely incorrect, but the fact that the method was trying to regulate pyruvate concentrations suggests that PYK might be the true target of regulation. Like the local MCA method, the RL method also regulated HEX1, GAPD and PGK.

In contrast, the unrestricted MCA method regulated the same reactions as in the low NADP/NADPH conditions, HEX1 and GAPD. The regulation under a high NADP/NADPH ratio is similar to the conditions in which NADP/NADPH is low primarily because increasing the NADP/NADPH ratio alone is insufficient to drive much flux through PPP. Because of less total regulation, the unrestricted MCA and RL methods result in significantly higher energy dissipation rates than the local MCA method and are thus likely to be more optimal regulation schemes.

### Regulation of PFK Maximizes Flux Through PPP

Increased flux can be channeled through the PPP if PFK activity is regulated to a greater extent or is turned off completely. Then significant flux flows through PPP instead of upper glycolysis and does so in a cyclical manner. There is experimental support for this as well. In *Neurospora crassa,* glycolysis and the PPP are circadianly regulated, with the PPP being regulated 180 degrees out of phase with upper glycolysis. In the extreme case when PFK activity is turned off in the model, then the cyclical operation of the PPP is such that three carbons are lost from each glucose molecule as CO_2_ before all the carbon reaches lower glycolysis as glyceraldehyde 3-phosphate.

In the case when PFK activity is set to zero, all methods apply regulation to HEX1. This is enough for the unrestricted MCA and RL methods to bring concentrations to within the observed experimental range, and both methods result in maximal energy dissipation rates (Figure 5). In contrast, the local MCA method additionally requires regulation in PPP at G6PDH, PGL and TKT. But even in this case, the local MCA method fails to completely bring sedoheptulose 7-phosphate into the range of the experimental observations. In attempting to control sedoheptulose 7-phosphate, the applied regulation is extensive enough such that the net flux through glycolysis, the pentose phosphate pathway and the TCA cycle approaches zero. Thus, the local MCA method fails to obtain control. In several cases involving the local MCA method, the concentration of sedoheptulose 7-phosphate and sometimes 6-phospho D-gluconate become uncontrollable resulting in concentrations higher than what is observed experimentally. The reason for this is that the respective reactions producing these compounds approach equilibrium; it is known that when a reaction approaches equilibrium, the concentrations of the products are no longer controllable^11^

In these cases, the reactions and their metabolites are effectively uncoupled from the non-equilibrium reactions. Lack of control may result in the respective metabolites reaching high concentrations in the cytoplasm, and the cytoplasm consequently becoming glassy and diffusion limited. Experiments support this principle. Recent reports provide evidence that active metabolism promotes cytoplasmic fluidization while inactive metabolism results in a glass-like cytoplasm with limited diffusion in both bacteria^10,12^ and eukaryotes^30^.

However, it is not clear that the failure to maintain control when using the local MCA method reflects poorly on the concept of modularity whereby enzymes use local control. The failure to obtain control of sedoheptulose 7-phosphate can also be due to the incomplete nature of the model of metabolism used here. It may be that in a more extensive model of metabolism, such as the inclusion of purine and pyrimidine biosynthesis pathways branching off of D-ribose 5-phosphate, control of sedoheptulose 7-phosphate by the local MCA method may be possible. We present this possibility because Transketolase (TKT), the enzyme producing sedoheptulose 7-phosphate is a key post-translational regulation point into purine synthesis^31^.

### Regulation Increases Reaction Free Energies

In all cases of regulation, whenever a reaction is regulated significantly, the reaction free energy is significantly different from the neighboring reactions which are not regulated, as shown in Figures 3B, 4B, and 5B. It is the act of regulating each reaction that causes the respective reactants to build up and products to become relatively depleted, which causes the free energy change of the reaction to increase in magnitude. This observation has wide support in the literature^13,14^, but the cause has been misinterpreted as being such that reactions are selected for regulation because they are far from equilibrium, rather than reactions being far from equilibrium because they are regulated.

**Figure 3.**
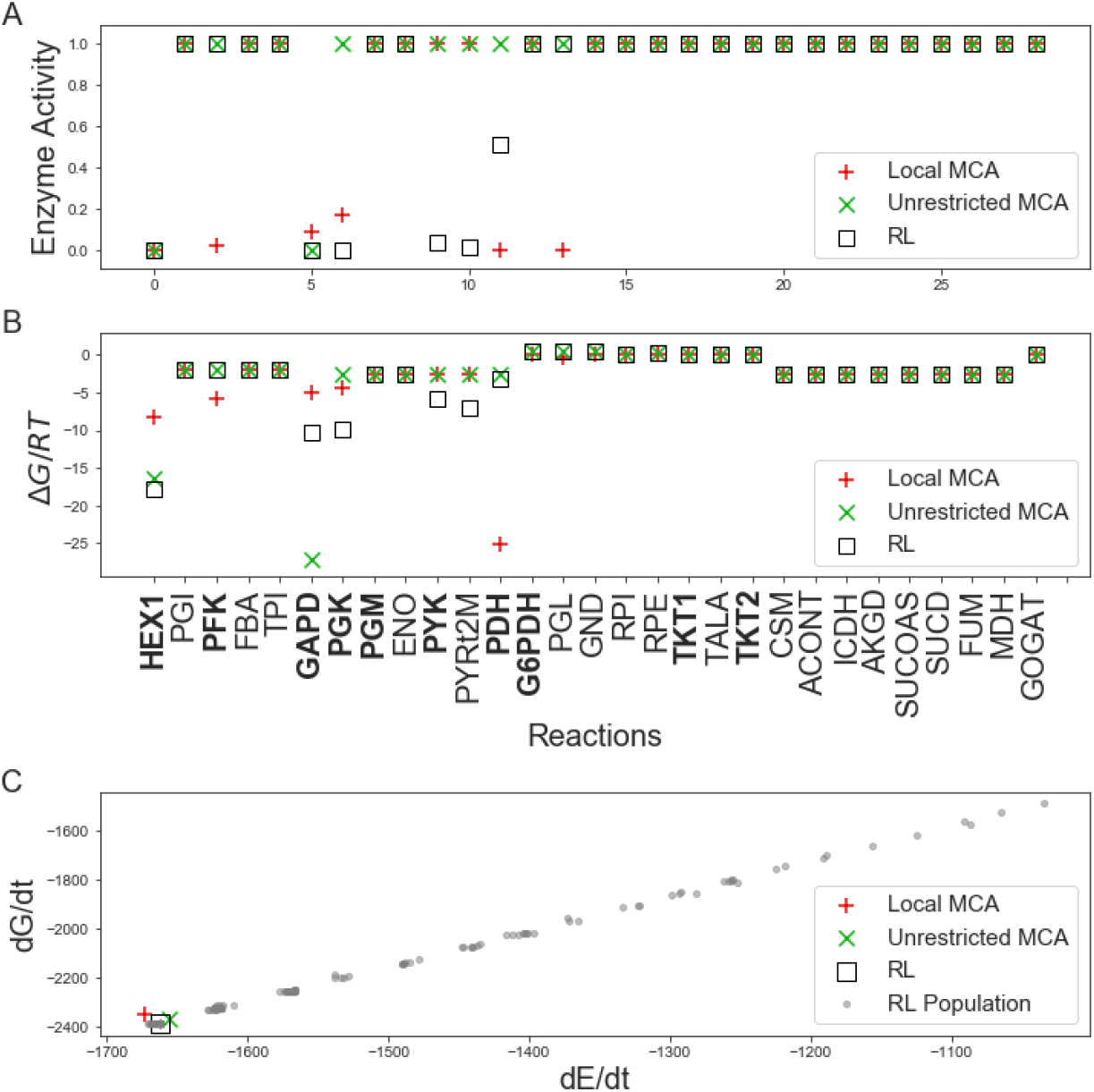
Glycolysis-PPP-TCA cycle predictions with high NAD/NADH and low NADP/NADPH conditions. (A) Predicted enzyme activities at terminal states are calculated using Metabolic Control Analysis, shown as red ‘plus’s and green ‘X’s, respectively. Results are compared to those found using a RL approach (black square).(B) Reaction free energy changes are no longer equally distributed across subpathways (Fig. 2, upper glycolysis, PPP, lower glycolysis, TCA cycle) but instead free energies are further from equilibrium at reactions where regulation is applied. (C) Free energy and energy dissipation rates. Grey dots represent the population of terminal states found while training the RL agent. **Abbreviations:** HEX1, Hexokinase; PGI, phosphoglucose isomerase; PFK, phosphofructokinase; TPI, Triosephosphate isomerase; GAPD, Glyceraldehyde 3-phosphate dehydrogenase; PGK, Phosphoglycerate kinase; PGM, phosphoglycerate mutase; ENO, Enolase; PYK, Pyruvate kinase; PYRt2m, pyruvate transporter; PDH, Pyruvate dehydrogenase; G6PDH, Glucose 6-phosphate dehydrogenase; PGL, Phosphogluconolactonase; GND, phosphogluconate dehydrogenase; RPI, Ribose 5-phosphate isomerase; RPE, Ribose 5-phosphate epimerase; TKT1, Transketolase 1; TALA, Transaldolase; TKT2, Transketolase 2; CS, Citrate Synthase; ACONT, Aconitase; ICDH, Isocitrate dehydrogenase; AKDG, a-ketoglutarate dehydrogenase; SUCOAS, Succinyl-CoA synthetase; SUCD1, Succinate dehydrogenase; FUM, Fumarase; MDH, Malate dehydrogenase; GOGAT, Glutamine oxoglutarate aminotransferase.

**Figure 4.**
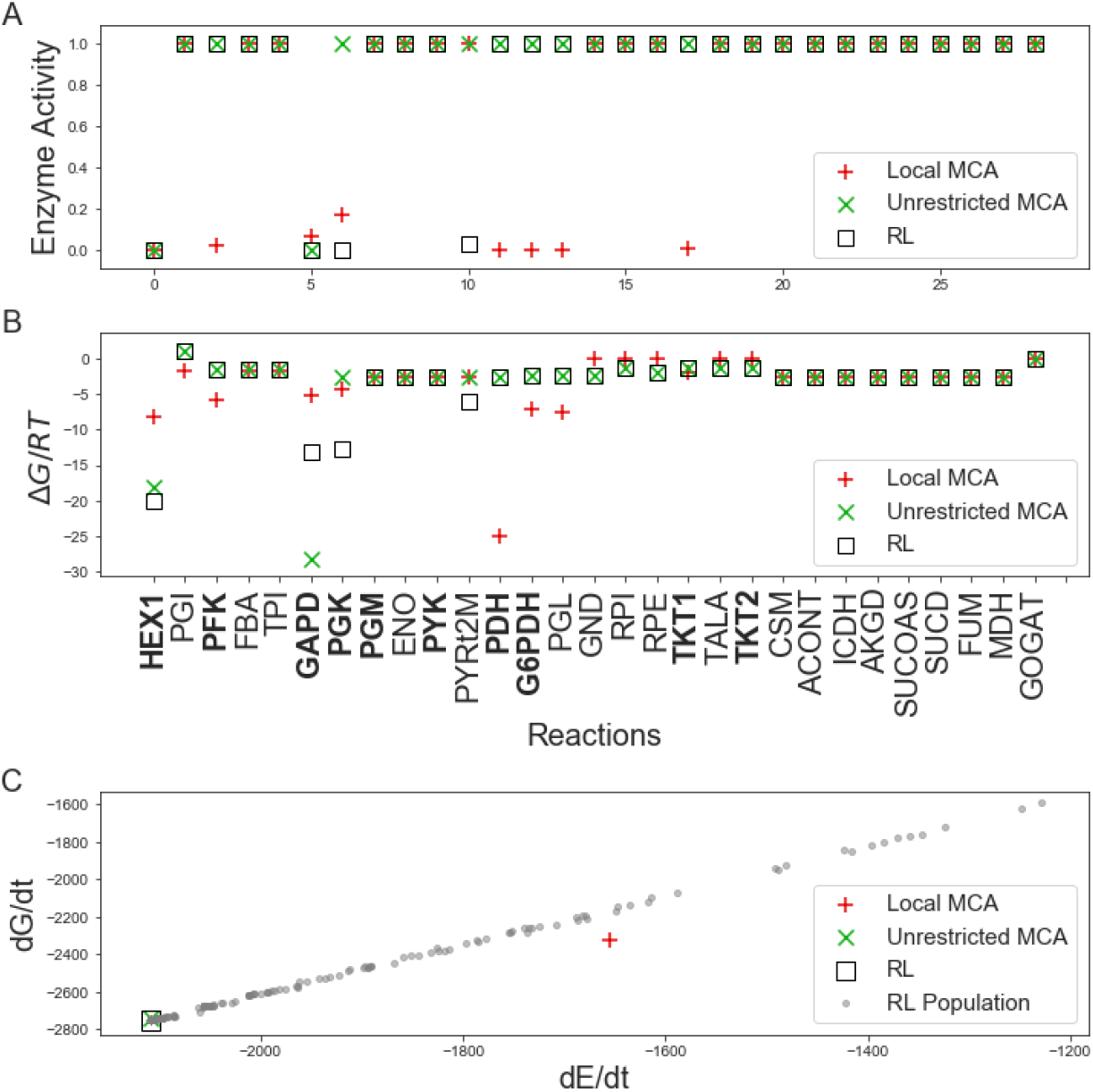
Glycolysis-PPP-TCA cycle predictions with high NAD/NADH and high NADP/NADPH conditions. (A) Predicted enzyme activities at terminal states are calculated using Metabolic Control Analysis, shown as red ‘plus’s and green ‘X’s, respectively. Results are compared to those found using a RL approach (black square). (B) Reaction free energies. (C) Free energy and energy dissipation rates. Grey dots represent the population of terminal states found while training the RL agent.

**Figure 5.**
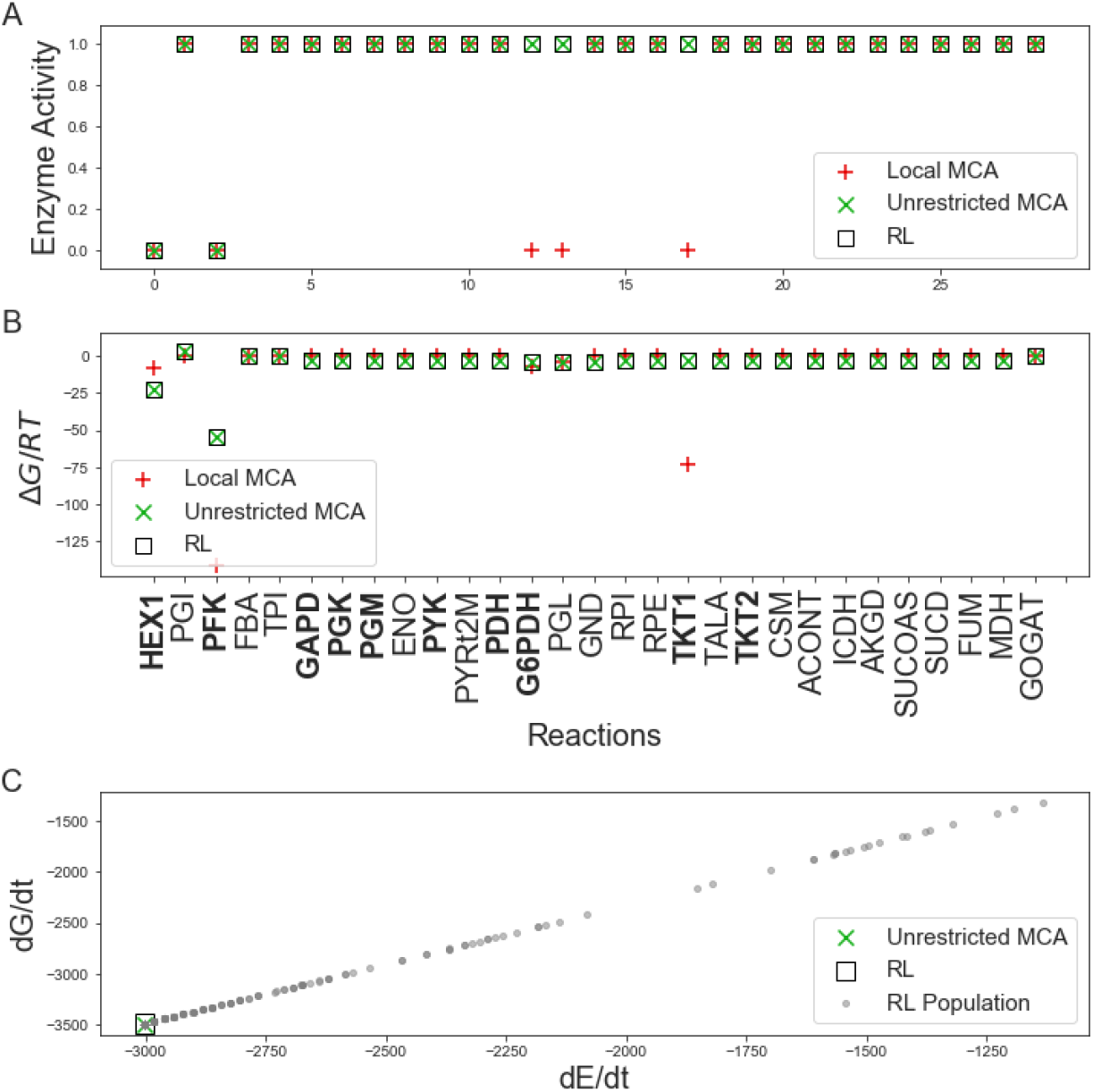
Glycolysis-PPP-TCA cycle predictions with high NAD/NADH and high NADP/NADPH conditions and PFK activity set to zero. (A) Predicted enzyme activities at terminal states are calculated using Metabolic Control Analysis, shown as red ‘plus’s and green ‘X’s, respectively. Results are compared to those found using a RL approach (black square). (B) Reaction free energies. (C) Free energy and energy dissipation rates. Grey dots represent the population of terminal states found while training the RL agent. The local MCA method results in zero flux (supplementary materials Table S1) and is therefore not shown.

The reasoning for assuming that highly non-equilibrium reactions are selected for regulation has to do with the established principle that biological systems activate metabolites for reactivity by covalently attaching high potential groups such as coenzyme A and phosphates. These reactions will then have much higher standard free energies of reaction than they would otherwise.

However, the use of such activators as phosphoryl groups and coenzyme A to drive a reaction will not just result in the respective reaction being further from equilibrium, but all reactions in the pathway will be further from equilibrium because increased product formation of the activated reaction will result in increased reactant concentration for the next reaction, and so forth, as the effect propagates down the pathway until a steady state is reached. As a result of the highly non-equilibrium nature of reactions in the pathway, many reaction products may be produced in biologically unreasonable concentrations. This problem is solved by reducing the activity of either the enzyme catalyzing the reaction or upstream enzymes that have control of the flow of material into the pathway. The reactions that have the most control can be determined using concentration control coefficients and thermodynamics.

## Discussion

All predicted schemes discussed above enforce regulation on enzymes that are known to be regulation sites. Nine of these 11 enzymes are known to be sites of post-translational regulation in glycolysis and the pentose phosphate pathway, either allosterically or through chemical modification (Table 1): hexokinase, phospho-fructokinase, glyceraldehyde phosphate dehydrogenase, phosphoglycerate kinase, pyruvate kinase, pyruvate dehydrogenase, glucose 6-phosphate dehydrogenase, transketolase, and pyruvate carboxylase (supplemental material). Only the pyruvate mitochondrial transporter (PYRt2m) and phosphogluconolactonase (PGL) are not known to be regulated. The regulation assigned to the pyruvate transporter was done stochastically by the reinforcement learning and likely should be assigned to PYK or PDH, as it was the deterministic MCA approaches. PGL presumably would be hard to control since it catalyzes a highly favorable ring opening which may only require desolvation in the enzyme active site. It is worth noting the enzymes that are known to be regulated but were not indicated as being regulated in this study. Foremost among these is fructose bisphosphatase (FBA), an enzyme that is well-known to be regulated in gluconeogenesis. Under the limited number of conditions used in the study of gluconeogenesis herein, levels of fructose 6-phosphate or other downstream products never rose high enough to require regulation. Likewise, the products of enolase, phosphoglucose isomerase, PEP carboxykinase, glucose 6-phosphatase never rose to the level that these needed to be regulated, but it would be reasonable to expect that the respective enzymes may need to be controlled under conditions that were not tested here.

**Table 1.** The set of enzymes found to be regulated in all analyses along with the associated pathway, the concentration control coefficient, 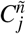, of the reaction summed over all metabolites before any regulation is applied, the method predicting the regulation and the experimental evidence from the literature for predicted regulation. Abbreviations are the same as in Figure 2. PC is pyruvate carboxylase and is observed to be regulated in gluconeogenesis (supplementary material Table S4).

| Enzyme | Pathway | $C_j^{\bar{n}}$ | Prediction Method | Evidence |
| --- | --- | --- | --- | --- |
| HEX1 | Upper glycolysis | 12.4 | RL, L-MCA, MCA | 13 |
| PFK | Upper glycolysis | 4.6 | L-MCA | 13,14 |
| GAPD | Lower glycolysis | 4.6 | RL, L-MCA, MCA | 14,43 |
| PGK | Lower glycolysis | 3.8 | RL, L-MCA | 14,44 |
| PYK | Lower glycolysis | 1.7 | RL | 45-47 |
| PYRt2m | Lower glycolysis | 1.1 | RL | — |
| PDH | Lower glycolysis | 0.6 | RL, L-MCA | 48 |
| G6PDH | Pentose phosphate | 16.8 | RL, L-MCA | 49 |
| PGL | Pentose phosphate | 16.0 | L-MCA | — |
| TKT | Pentose phosphate | 5.0 | L-MCA | 31 |
| PC | Gluconeogenesis | 3.7 | RL, L-MCA, MCA | 50 |
| **Modeled and known to be regulated but not observed** |  |  |  |  |
| PGM | Lower glycolysis | 3.0 | — | 51 |
| FBP | Gluconeogenesis | 0.1 | — | 13,14 |

Of the 11 enzymes predicted to be regulated, outsized roles were played by the branch points of each of the pathways, as quantified by the influence of the enzyme activity coefficients, 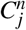, on the respective reactants or products (Table 1). The summary concentration control coefficient reports the total influence of the activity of the enzyme on all metabolites exceeding the experimentally observed values. Hexokinase, the entry point into the model and entry point into upper glycolysis and the pentose phosphate pathway, had the largest role with 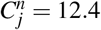, meaning that hexokinase effectively had 100% control over 12.4 reactions. Likewise, glucose 6-phosphate dehydrogenase, the entry point into the PPP, had effectively 100% influence over 16.8 reactions, although this value is only seen this high when the phosphofructokinase activity is set to 0.0 such that the PPP acts cyclically and three circuits around the cycle are made for each glucose metabolized. Likewise, for lower glycolysis the main control point, glyceraldehyde 3-phosphate dehydrogenase, is the entry into the pathway which is also where upper glycolysis and PPP converge. No regulation was needed for the TCA cycle under the conditions studied.

While the predictions align well with known sites of post-translational regulation, the predictions offer no information on whether the regulation would be due to allosteric interactions or chemical modification as might be inferred from more complex and expensive approaches that utilize (and require) absolute metabolite concentrations, fluxes inferred from isotope labeling studies, MS proteomics analyses and detailed kinetic models that include explicit enzyme binding, catalysis and product release^2^. The regulation predictions provided here, however, were done purely *in silico* with the optional use of absolute metabolite concentrations, if available. Although the regulatory effector can’t yet be inferred from this approach, it would seem reasonable to assume that control of metabolite concentrations would be due to allosteric regulation since allosteric interactions work on a faster time scale than post-translational modifications. It is likely that post-translational modifications act to redirect flux when either degradation of enzyme would be too slow, or when degradation and later resynthesis of the enzyme would be too costly^32^, which is not the scenario addressed here.

Both MCA approaches were based only on adjusting the activities of enzymes that would have the most influence on reducing concentrations to physiological values. Only the RL approach rewards regulation schemes for maximizing the entropy production rate (Eqn. (26)). Even though the RL and MCA methods have different aims, both maximized the energy dissipation rate, *dE/dt*, a principle alluded to by Lotka^33^. Furthermore, while the unrestricted MCA approach and the RL performed similarly, the local MCA approach did not always find a solution, which could reflect the incompleteness of the metabolic network that is modeled, or may simply indicate that modular regulation to this degree is insufficient. In addition, in at least one case the local MCA approach did not produce solutions with the highest energy dissipation rates. However, the set of enzymes predicted by the local MCA approach covers many more of the enzymes known to be classically regulated, as shown in Table 1.

Consequently, we have shown how post-translational regulation results in the emergence of the general principle of maximal, entropy production rate for metabolism, and we can now also include the principle of maximization of the energy production rate, *dE/dt*, for pseudo-steady state phenotypes. When these principles are applied for predictions, each prediction must include the physicochemical constraints on the system, such as the inherent constraints on the maximal rates of enzymes and thermodynamic costs and benefits, not simply metabolite solubilities^32^. These additional physicochemical constraints can explain the observed upper limit to free energy dissipation in microbial systems^34^.

The observation of an upper limit to free energy dissipation is related to the concept of maintaining the adenylate energy charge ratio. The adenylate energy charge rule widely found in textbooks was defined in terms of concentrations as [(ATP) + 0.5 (ADP)]/ [(ATP) + (ADP) + (AMP)]. It was proposed that biological systems maintain values of the energy charge between 0.75 and 0.90. There are now many known exceptions to this proposed rule that it can no longer be regarded as a rule but as an emergent property, just as the maximization of energy production rates is an emergent property due to natural selection.

The simulation-based predictions of enzyme activities presented in this paper advance both the practice and theory of biology. The ability to predict from simulation or infer the free energy changes and control coefficients (in addition to fluxes) for each reaction allows the use of control theory and learning to analyze and explore the operations of the cell. In synthetic biology the development of cell lines often requires additional circuits and can result in unforeseen consequences or lower cell growth rates. Simulation of cells with engineered or deleted circuits will allow prediction of the effects in place of difficult trial and error in experiments.

Finally, it is important to understand the principles behind post-translational regulation because regulation of metabolism is precisely what controls a cell’s energetic behavior. From bacterial growth and reproduction, to developing cells or even halting the growth of cancer cells, regulation plays the central role. Learning how cells regulate and control themselves is essential for designing new organisms that have an intended purpose (synthetic biology), developing new strategies to target and control microbial and metabolic diseases (medicine), and understanding design principles of biology (fundamental science). Currently no other experimental or computational approach has been shown to identify points of regulation in metabolism in a rapid manner.

## Methods

### Convex Optimization Approach for obtaining Metabolic Steady State

For a reversible chemical reaction, the reaction is described by,

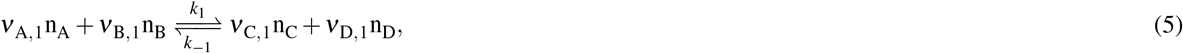

where *A, B, C, D* represents the molecular species, the concentrations are given by *n_i_, i* = {*A, B, C, D*}, and *v_i,j_* represent the unsigned stoichiometric coefficients for each molecular species *i* in the forward and reverse reactions *j* = {1, −1}.

The law of mass action may be formulated in terms of chemical kinetics or thermodynamics. With respect to chemical kinetics, the law of mass action is expressed by the rate or net flux, *J*_*net*,1_, of the reaction where the forward and reverse rates are proportional to the respective reactants,

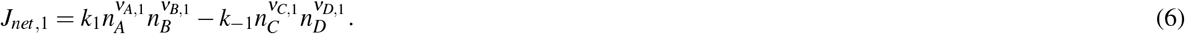

In this formulation, *k*_1_ and *k*_-1_ represent the rate constants of the forward and the reverse reaction, respectively. On the other hand, the thermodynamic expression of the reaction utilizes the change in free energy, *G*, with respect to the extent of a reaction, *ξ*. The ratio of the respective reactants and products are combined to form the reaction affinity, *A*_1_ = *∂G*/*∂ξ*_1_, such that,

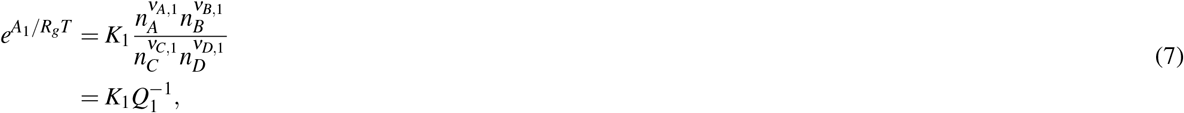

where *K*_1_ = *k*_1_/*k*_-1_ is the equilibrium constant and *Q*_1_ is the reaction quotient. Also, the analogous equation for the reverse reaction is the reciprocal,

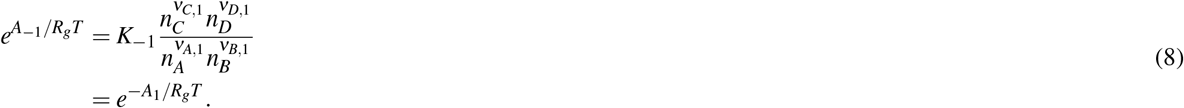

Note that Eqn. (6) is a purely kinetic description of the law of mass action, while Eqns. (7) and (8) are purely thermodynamic expressions. This results from the fact that the latter equations do not contain any information on the time dependence of the reaction. These formulations, however, are not mutually exclusive. Time dependence and thermodynamics can both be described in a single equation by factoring the opposing rate from each term of Eqn. (6),

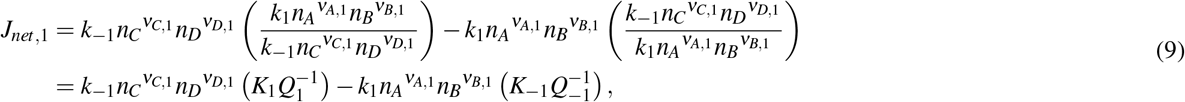

where *K*_1_ and *K*_-1_ are the equilibrium constants and *Q*_1_ and *Q*_-1_ are the reaction quotients for reaction 1 and −1, respectively. Eqn. (9) is the Marcelin-de Donder equation^15,35^, which describes the forward and reverse reactions as being functions of the time independent odds of the reaction and the rate of change of the odds.

Given a metabolic model with *Z* reactions, *M* metabolic species, and *N* total particles, we formulate the flux through each reaction using Eqn. (9). In this work, the largest values of *Z* and *M* in a pathway are 29 and 47 respectively. If we assume the rate of change of the odds are equal and independent of concentrations, then the coupled reactions occur on the same time scale. Under these assumptions, the resulting equation for the *j^th^* reaction is the Marcelin equation^36^,

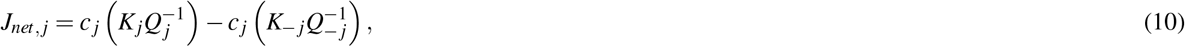

where *c_j_* represents the time dependence of the reaction odds. Because the exponential family of distributions are always log-concave when counts are greater than or equal to zero, the energy surface on which the reactions occur is convex. This is achieved by expressing the reactions as functions of the reaction affinities via Eqns. (7) and (8),

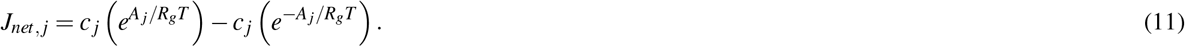

A vector of *Z* reaction fluxes *J* = [*J*_1_,…, *J_Z_*]^*T*^ can be determined from the *M* by *Z* stoichiometric matrix *S* and the *M* chemical potentials. The stoichiometric matrix consists of elements *γ_i,j_*, which are the signed stoichiometric coefficients for chemical species *j* in reaction *i*. The time dependence of the vector of counts *n* = [*n*_1_,…, *n_M_*]^*T*^ of chemical species is,

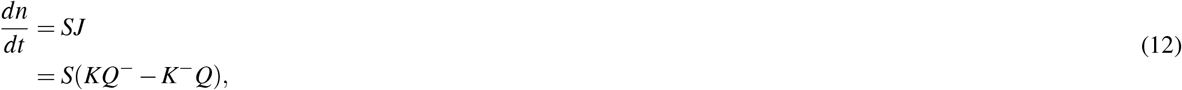

where *SJ* is the matrix multiplication between *S* and 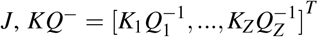 is the vector of thermodynamic odds for the forward reactions, and 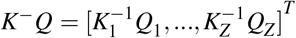 is the vector of thermodynamic odds for the reverse reactions. Without any constraints applied, Eqn. (12) will converge to an equilibrium solution, whether the equation is solved using ordinary differential equations or optimization methods. To obtain a non-equilibrium steady state, non-equilibrium boundary conditions must be applied. In this case, the non-equilibrium boundary conditions consist of boundary metabolite values representing the reactants and products of the overall chemical process that are held fixed. If there are *M_V_* variable species and *M_B_* = *M – M_V_* boundary (fixed) species such that 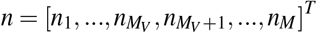, then the stoichiometric matrix will contain a non-singular submatrix and Eqn. (12) will have unique solutions only if *M_V_* ≤ *Z*. The vector of counts *n* can be split into subvectors 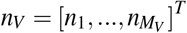 and 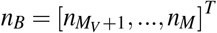 such that 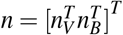. Likewise, the stoichiometric matrix can also be split along the rows representing metabolites to separate the entries for the variable metabolites from those for the boundary metabolites such that 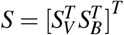 where *S_V_* is an *M_V_* by *Z* matrix and *S_B_* is *M_B_* by *Z*. The time dependence of each of the chemical species is given by,

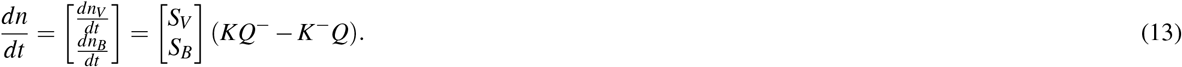

The optimization problem is to find *n_V_* satisfying

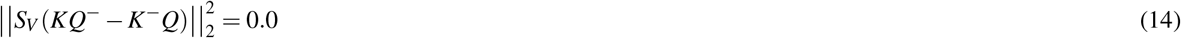

subject to the *M_B_* boundary conditions. The optimization is carried out with a nonlinear least-squares approach using the Levenberg-Marquardt method^37,38^, and solves for the concentrations of the chemical species which makes up the reaction quotient, *Q*. When *S_V_*(*KQ*^-^ – *K*^-^*Q*) = 0.0, the optimization has found a kinetic steady state as well as a thermodynamically balanced state such that the net thermodynamic driving forces on all the reactions are equal for linear pathways, or for branched pathways, the net thermodynamic driving forces are proportional to the stoichiometry. If one is only interested in the thermodynamic properties, fluxes and concentrations at steady state, then there is no need to solve for the rate constants. Otherwise, rate constants can be back-calculated and used to solve for the system dynamics using, for example, Eqn. (6). For example, setting *j* = 1, Eqn. (9) can be solved for *k*_±1_ as follows:

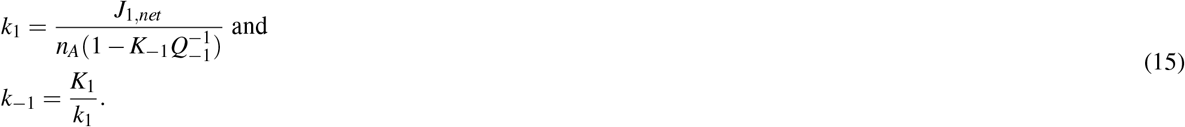

The kinetically accessible energy surface is not necessarily convex because of the introduction of the rate constants - each reaction now has its own time dependence.

The predicted concentrations from the optimization follow the multinomial Boltzmann distribution in which the concentration of each species is proportional to its standard chemical potential, 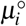, adjusted for aqueous solution at pH 7.0,

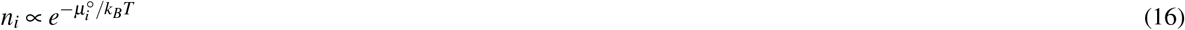

subject to the constraints of the reaction stoichiometries and the non-equilibrium boundary conditions. The boundary conditions consist of fixed concentrations of environmental nutrients such as glucose and waste products such as CO_2_, as well as some cofactors. Because the concentrations are distributed as a function of their standard chemical potentials in aqueous solution, the concentrations of highly hydrophilic charged species may be orders of magnitude above physiological values. For instance, concentrations of ATP or acetyl CoA may be on the order of ten molar or more. Such high concentrations would make the cytoplasm highly viscous such that diffusion would be slowed down significantly, and cellular activity would come to a halt. However, as we shall show, the concentrations can be brought into alignment with physiological values using enzyme activities determined from Metabolic Control Analysis^39,40^.

### Metabolic Regulation: A Metabolic Control Theory Approach

Regulation is applied to reactions by changing the scalar valued activity of the *j^th^* enzyme, *α_j_* ∈ [0.0,1.0], where activity values of 0.0 and 1.0 represent complete reaction regulation and no enzyme regulation, respectively. The activity for each reaction *j* is represented by a multiplier to the net reaction flux *J_j_* such that,

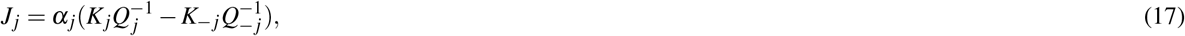

and likewise,

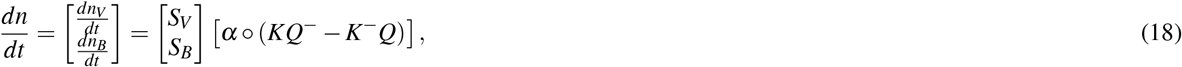

where ○ represents the Hadamard element-wise product. Since any reaction may be regulated, the state of the system can be described by the activity vector, *α*, steady state fluxes, *J*, and steady state metabolite concentrations *n*. Because the latter two state variables can be determined from a fixed set of activities via the optimization routine, system states can be defined simply by the activity vector *α* instead of the tuple (*α, J, n*).

In Metabolic Control Analysis (MCA), the sensitivity of a concentration *n_i_*, to the activity *α_j_* of enzyme *j* is defined as the concentration control coefficient 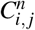 (Eqn. (1)). Concentration control coefficients can be used to determine how much to reduce the activities of an enzyme to bring the predicted concentrations into alignment with physiological values observed from experimental metabolomics assays. The detailed calculation is described in the supplementary material. If concentrations *n_i_* for a metabolite *i* have not been measured, then target values are assumed to be 1.0 millimolar, which ensures that concentrations stay reasonable even for metabolites whose concentrations have not been measured. When predicted values exceed the measured or target values, regulation is applied to reactions by changing the scalar valued activity of the *j^th^* enzyme, *α_j_*.

Which reaction to regulate is determined from examining the concentration control coefficients with regard to the metabolites whose concentrations are higher than is observed in experiment. We denote the set of such metabolites as *M*′ = {*i*|*ñ_i_*, > *n_i_*}. An activity is then selected to be reduced based on the influence that the activity has on these concentrations,

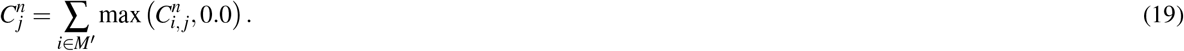

Because activities are reduced from initial values of 1.0 (full activity), only 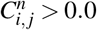 are considered in the sum so that reduction in activity correlates with reduction in concentration. A component cost function, *L_i_*, is defined as the division of the predicted concentrations or counts to the measured concentrations or counts, *L_i_*, = log(*ñ_i_*/*n_i_*). In order to determine the point where steady state metabolite levels are ‘in caliber’, we utilize a stopping criteria function that returns a positive scalar if any *L_i_*, < 0.0 and returns zero once *L_i_*, ≤ 0.0 for all *i*. We define this cost function as follows:

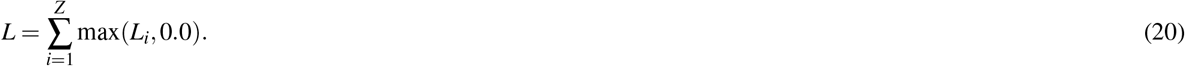

The maximum of *Li* or zero is used because the model only predicts metabolite populations that are free in solution, but the experimentally measured concentrations are in principle those that are both enzyme-bound and free in solution. Thus, concentrations from predictions are assumed to be ‘in caliber’ with experimental data if the predicted concentrations are less than or equal to experimentally measured concentrations (*L_i_*, ≤ 0.0).

In practice, the activity that reduces the cost function, *L*, the greatest amount is chosen for regulation and is again determined using MCA. In MCA, the concentration control coefficient for metabolite *i* due to control by reaction *j* is defined by Eqn. (1). Consequently, the change in concentration or counts due to a change in activity of reaction *j* is,

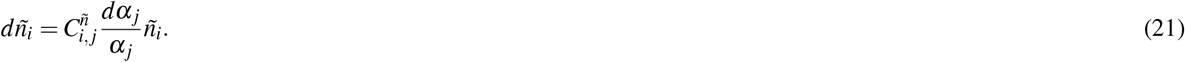

For metabolite *i* with predicted concentration *ñ_i_*, and a target concentration of *n_i_*, the estimated change in the costs, Δ*L_i,j_*, of metabolite due to a change in activity *α_j_* of reaction *j* is:

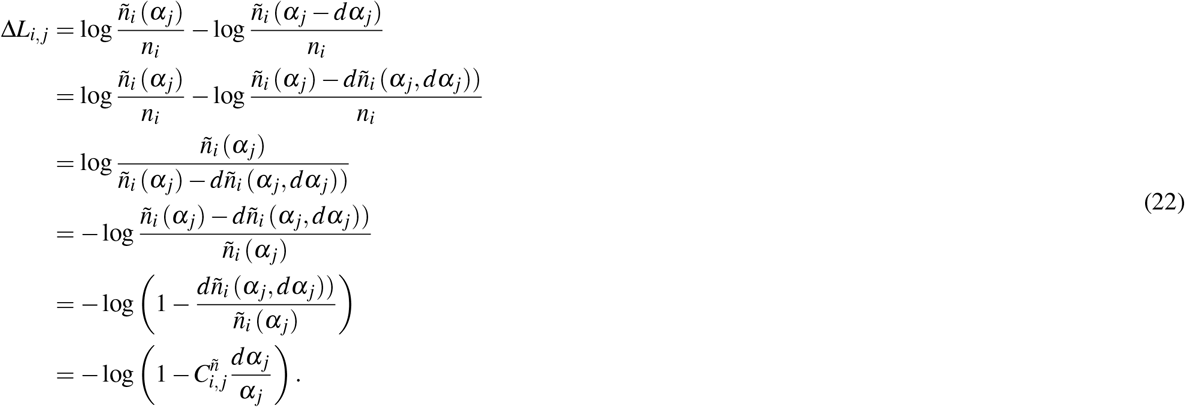

The change in total costs over all metabolites due to a change in activity of reaction *j* is calculated by summing over metabolites that are out of ‘caliber’ with respect to the experimentally observed concentrations. We calculate the total cost as follows:

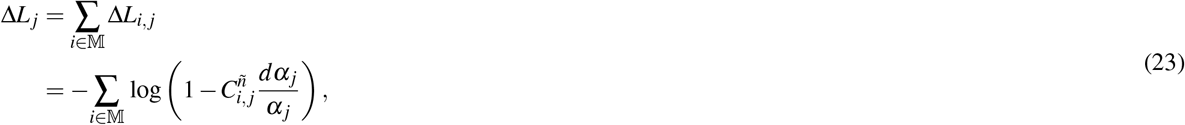

where 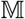 represents the set of reactions able to be regulated or controlled. Finally, the question of which enzymes should be allowed to be control points must be addressed. Two approaches were taken with MCA: an unrestricted control approach in which any enzyme could be a regulator for any metabolite, and a restricted approach in which only enzymes whose immediate products exceeded the target values could be considered as a regulator. We refer to the latter as a local-control approach (MCA Local) since an enzyme’s immediate products (and possibly other metabolites) are being controlled. Regulation is then applied at the reaction maximizing,

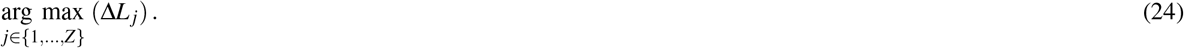

Once a reaction *j* is chosen, the activity *α_j_* is changed by an appropriate amount (supplemental material). When all metabolite values are brought into agreement with experimental observations, rate constants can be determined, if desired, using Eqn. (15). Alternately, the influence of the activities can directly be incorporated into the rate constants. For example, given *j* = 1, the resulting rate constant is,

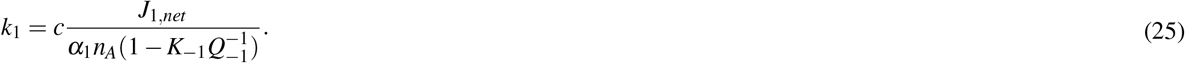

However, there is an important conceptual difference between solving mass action rate laws with parameters based on the approach provided by Eqn. (15) compared to Eqn. (25). While the former assumes regulation is needed to bring concentrations under control, the latter assumes no regulation is needed and control is hardwired into rate constants. The advantages of the former are two-fold: (1) under different nutrient conditions, enzyme activities can be altered to control metabolite concentrations; and (2) enzyme activities are adjusted away from the maximal entropy distribution only enough to bring concentrations into alignment with observed values, resulting in a more favorable total free energy of the system. A lower total free energy also would reduce the cost of replicating of metabolism. The actual balance between these two approaches will likely be a middle ground between the reliance on activity coefficients as opposed to rate constants. It is unlikely that enzymes can evolve such that the ideal rate constants, i.e. those implied by Eqn. 15, are possible for every reaction. Instead, rate constant values will be limited by constraints due to the physics of the catalytic process.

### Exploring Regulation: A Reinforcement Learning Approach

The MCA method for bringing the predicted concentrations in alignment with observed concentrations is a deterministic approach based on an assumption that metabolite concentrations depend linearly on the enzyme activities. It is feasible that the assumption of linearity used in the MCA analysis (supplemental material) results in sub-optimal regulation. Optimal regulation has been hypothesized, based on empirical data, as regulation that maintains a high energy charge, defined in terms of ATP, ADP and AMP^5^. A less *ad hoc* definition of optimal regulation would be the maximization of the entropy production rate, which has also long been hypothesized as an objective of biological systems^33,41^. Neither of these concepts are addressed in the MCA approaches discussed above. For steady state systems, the entropy production rate (EPR) is the negative of the free energy dissipation rate^27,28^,

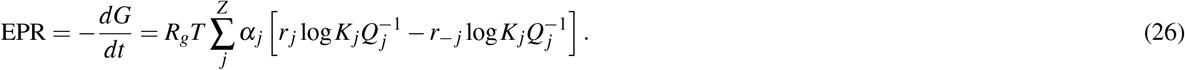

Given a goal of maximizing the EPR, it is not clear which MCA protocol above, if either, would maximize the entropy production rate. On one hand, the unrestricted MCA method uses less regulation and therefore often results in higher reaction fluxes, which would increase the EPR (Eqn. (26)). On the other hand, entropy is maximized when the value of the argument of the logarithms are distributed as uniformly as possible, which is the opposite of what occurs when a minimal set of enzymes are chosen to be regulated. In order to explore the regulation space more completely to investigate these issues, we utilize a machine learning method that avoids the linearity assumption by directly testing multiple future states and is directly rewarded for maximizing the EPR.

Specifically, we use a Reinforcement Learning (RL) framework which can address decision problems that are otherwise combinatorially intractable. Even a small metabolic network may have on the order of 20-50 reactions. To explore the state space fully using the deterministic MCA approach, on the order of 100-500 decisions need to be made as to which reaction to regulate depending on the state of the system. The search space is then approximately between 20^100^ and 50^500^, a number much too large to tackle by an exhaustive search or Monte Carlo approach.

In our framework, optimal regulation of a metabolic network requires that the EPR be maximized while satisfying a stopping criteria: *L* = 0.0. A diverse set of reaction regulation schemes represented by enzyme activity values, {*α*_1_,…, *α_Z_*}, satisfy the stopping criteria, but each scheme results in a different EPR (Fig. 3C–5C, grey dots). Thus, we utilize a hybrid optimization-RL approach to iteratively search for the best regulation scheme. (A hybrid simulation-RL approach can also be used.) In this framework, the agent iteratively learns how to navigate a state space, *S*, by using different possible actions from an action space, *A*. States correspond to the value of enzyme activities while actions correspond to regulating a specified reaction. We therefore define the state space *S* as the subset of ℝ^*Z*^ using range of each enzyme activity, [0.0,1.0]^*Z*^, and the action space as the set of reactions, *A* = {1,2,…,*Z*}. We also define a subset of *S* where learning terminates, *S_T_* = {*s* ∈ *S*|*L*(*s*) = 0.0}.

Because regulating the *j^th^* enzyme results in a deterministic step-size, Δ*α_j_*, the resulting state is given by the following set of enzyme activities: { *α*_1_,…, *α_j_* – Δ*α_j_*,…, *α_Z_*}. The goal of Reinforcement Learning is to learn an optimal policy, *π* *: *S* → *A*, which results in a regulation scheme that maximize some defined notion of rewards, *R*: *S* × *A* × *S* → ℝ. In other words, learning the optimal policy corresponds to learning the regulation scheme for the chemical reaction network that results in the largest reward.

Each reaction that is regulated results in a scalar valued reward, or feedback, from the environment based on an action/regulation (Figure 1) that indirectly defines optimal regulation schemes. Each regulation decision alters the steady state metabolite concentrations, which are obtained from optimization or simulation of Eqn. (18), and used to calculate rewards using a loss function, Λ, specified by

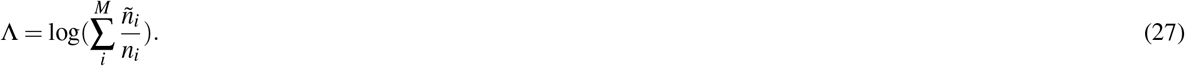

The formulation of Λ emphasizes regulation of reactions that affect metabolites which are furthest from being in caliber with experimental measurements.

We define the environmental feedback, or reward function *R* as:

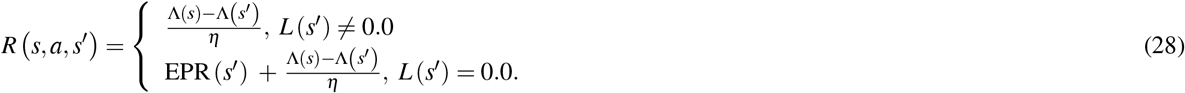

Intermediate rewards are calculated by the reduction in Λ scaled by a positive factor *η*. Once a terminal state is found, the final reward consists of the final change in the scaled loss function as well as the entropy production rate calculated at the final state, 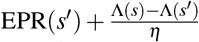. Thus, the agent aims to both increase the value of EPR(*s*) for *s_t_* ∈ *S_T_* while satisfying the constraint *L* = 0.0 and regulating as many reactions as is necessary.

Learning is conducted by iteratively updating the current policy function, *π*: *S* → *A*, that determines the agent behavior. The policy function determines which reaction *j ∈ Z* should be regulated based on the current enzyme activities, {*α*_1_,…, *α_Z_*} ∈ *S*. Here, we utilize an n-step SARSA algorithm^25^ to perform fitted value function iteration. An optimal policy is therefore learned by iteratively updating the value function, *V*: *S* → ℝ, which is defined as the expected rewards to be received by following a fixed policy from a specified state, 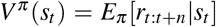. In an n-step algorithm, the value function is meant to predict the discounted reward, *r_t:t+n_*, for n future steps. The n-step reward experienced by the agent is defined as 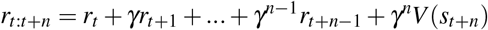, where *γ* ∈ [0.0,1.0] is the discount factor. Each reward, *r_t_* = *R*(*s_t-1_,a,s_t_*), represents the feedback from moving into state *s_t_* from *s_t-1_* after taking some action *a*. The first n steps represent the rewards experienced, while the term *V*(*s_t+n_*) represents the future rewards. Once a terminal state is less than *n* steps away, the n-step reward is truncated to the appropriate length.

Learning the value function implicitly improves the policy. The relation between the value of a state and the policy is given by an *ε*-greedy policy, which is defined as:

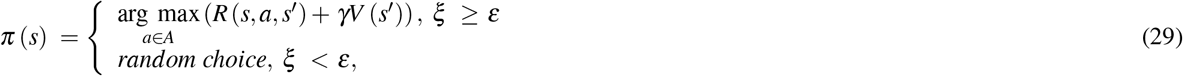

where *ξ* is a uniform random number between 0.0 and 1.0. As the value function is better estimated, the policy determines reactions to regulate that lead to the greatest cumulative reward. Exploration is imposed by randomly choosing reactions to regulate, allowing the policy to escape local minima. As the agent learns, *ε* is slowly annealed to reduce exploration and fluctuations in the value function. During each training episode, we begin at the state *s* = {1.0,…, 1.0}, such that all enzyme activities are unregulated. Trajectories through state space are stopped when the stopping criteria *L* = 0.0 is satisfied. This condition requires that all reactions have cost function values at or below zero before the reinforcement learning ends and the predictions are in caliber with the experimental values.

Finally, the state value function is estimated by using a neural network implemented in PyTorch^42^ with a single hidden layer and hyperbolic tangent activation functions. Updates to the value function are performed by optimizing the neural network using stochastic gradient descent. This is done by backpropagating the squared loss between the predicted value and the n-step reward, [*V*(*s_t_*) – *r_t:t+n_*]^2^.

### Model Training

Prediction of network regulation was performed using a trained neural network to estimate the value function. Network weights were adjusted using stochastic gradient descent with a learning rate, *lr* ∈ {10^-4^, 10^-5^,10^-6^}. Each algorithm learned and generated data using an *ε*-greedy policy with initial *ε* = 0.5 or 0.2 depending on the size of the pathway. *ε* was annealed by dividing by a factor of two every 25 learning episodes.

For each pathway, 10 agents are trained for each different value of *n* ∈ {2,4,…, 12} and each learning rate. The resulting average of 10 RL runs for the glycolysis-PPP-TCA pathway (Figure S1) show the mean reward for the 350 training episodes. Optimal regulation is prescribed by analyzing the agent with the largest cumulative reward averaged over the last 50 terminal states.

### Experimental Data

The metabolomics data used in this study was from *E. coli* studies by Bennett, et al.^18^, and Park, et al^19^. Briefly, *E. coli* cells were grown in isotope-labeled media and then extracted in organic solvent containing unlabeled internal standards in known concentrations. Metabolites were extracted in cold solvent and analyzed using chromatography-MS, and concentrations relative to the known standard concentrations were obtained using peak ratios of the labeled samples to unlabeled standards.

## Acknowledgements

S. Britton was supported by a U. S. Department of Energy (DOE), Office of Science Graduate Student Research award and by funding from the DOE Office of Biological and Environmental Research, through project 74860. W. R. Cannon was supported by joint funding from the National Institute of Biomedical Imaging and Bioengineering through award U01EB022546 and by the DOE Office of Biological and Environmental Research, through project 69513. PNNL is operated by Battelle for the US Department of Energy under Contract DE-AC06-76RLO. M. Alber was supported by funding from the National Science Foundation, Grant DMS-1762063, through the joint NSF DMS/NIH NIGMS Initiative to Support Research at the Interface of the Biological and Mathematical Sciences.

## Supplementary information (optional)

See PDF file supplementary material.

