## Supplemental Material for "Machine Learning and Optimal Control of Enzyme Activities to Preserve Solvent Capacity in the Cell"

#### Supplementary Materials

##### Model training

Reinforcement learning agents are trained by iteratively learning the value function of each state, $s_t$. The value at a state represents the expected reward to be achieved from following the current policy: $V(s_t) = E[r_{t:t+n}|s_t]$. At each state with $t \geq n$ the squared error between the value of the state, $V(s_t)$, and the experienced rewards, $r_{t:t+n}$, is back-propagated to calculate appropriate changes in the neural network weights. As agents explore different possible regulation schemes, rewards are accumulated and averaged over each episode of training. Average rewards per episode are shown over the 350 training episodes for the gluconeogenesis and glycolysis-TCA pathways (Figure S1A) as well as the glycolysis-PPP-TCA pathway for each of the environmental conditions (Figure S1B).

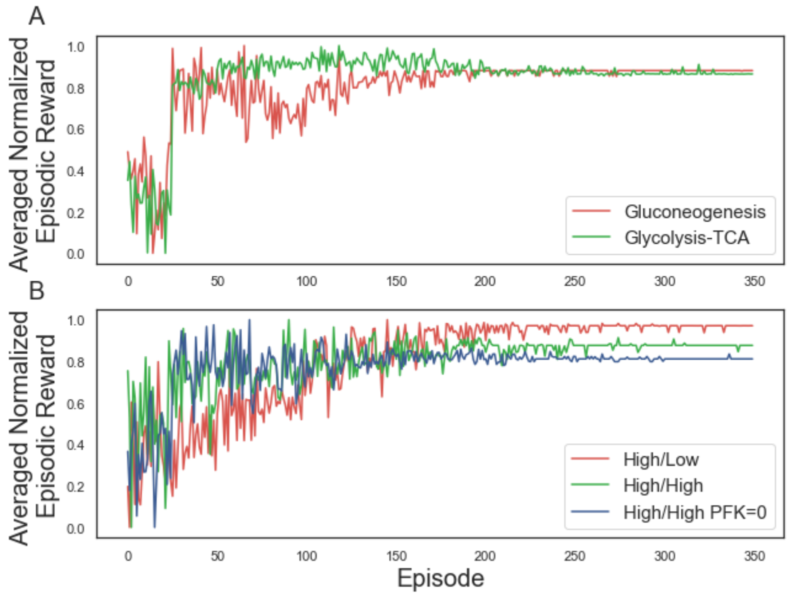


**Figure S1.** Cumulative normalized rewards averaged over 10 RL runs for the hyper-parameters (n, lr) which resulted in the maximal reward.

##### Calculating concentration control coefficients

Enzyme activities begin from a value of 1.0, i.e. the enzyme is unregulated. The current value of the activity is adjusted using Metabolic Control Analysis (MCA)[1]. In MCA, the concentration control coefficient is found by first computing the $M$ by $M$ symmetric linear stability matrix, $A^{n^*}$, given by,

$$A^{n^*}_{ij} = n^*_j \sum_{k=1}^{Z} S_{ik} \frac{\partial J_k}{\partial n_j}|_{n=n*}. \quad (1)$$

Here $n^*_i$ is the current concentration of metabolite $i$, $S$ is the stoichiometric matrix and $J$ is the vector of reaction fluxes. The concentration control coefficient for metabolite $i$ due to reaction $j$ is then,

$$\begin{aligned} C^n_{i,j} &= \frac{\partial \log n_i}{\partial \log \alpha_j} \\ &= \frac{\alpha_j}{n^*_i} \frac{\Delta n_i}{\Delta \alpha_j} \\ &= -(BS)_{ij} J_j, \end{aligned} \quad (2)$$

where $B = A^{-1}$. Note that the calculation of $C^n_{i,j}$ assumes metabolite concentrations are linearly dependent on enzyme activities. This assumption can be used to isolate the change in activity, $j$, needed to make a change in the product concentration $n_i$:

$$\Delta \alpha_j = \alpha_j \frac{\Delta n_i}{n^*_i} \left( - \sum_{k=1}^{Z} B_{ik} S_{kj} J_j \right). \quad (3)$$

In practice, when $C^n_{i,j} >> 0.0$ the assumption of a small change $n_i$ used in MCA is no longer valid and instead the current activity $j$ is instead updated using $\alpha_{j,new} = \alpha_{j,current}/5$. As the cost function $L$ (Methods Eqn. 20) approaches zero, then Eqn. 3 can be applied.

##### Analysis of gluconeogenesis pathway

The gluconeogenesis pathway is analyzed at low NAD/NADH ratio (0.02). The pathway has two known regulation sites fructose 1,6-bisphosphatase (FBP) and pyruvate carboxylase (PC). While both can be utilized to bring steady state metabolite concentrations into agreement with experimentally observed values, regulation of pyruvate carboxylase results in a larger energy dissipation rate ($dE/dt$). Regulation of alternative enzymes results in lower flux through the pathway and less energy available for use. Optimal predicted enzyme activities are in agreement for each method (Figure S2A). Table S4 lists the complete reaction activity, flux and free energy for each respective prediction method.

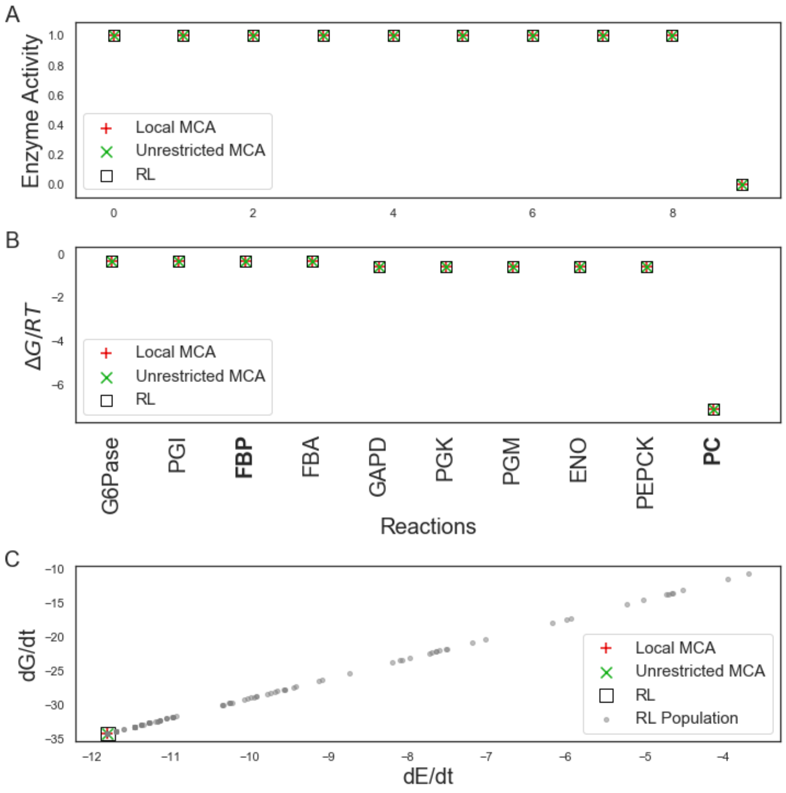


**Figure S2.** Gluconeogenesis cycle predictions with low NAD/NADH initial conditions. Predicted enzyme activities (A) and free energy (B) at terminal states are calculated using concentration control theory, shown as red 'plus's and green 'X's, respectively. Results are compared to those found using a RL approach (black square). Grey dots (C) represent the population of terminal states found while training the RL agent.

##### Analysis of glycolysis-TCA pathway

The glycolysis-TCA pathway is a subset of the larger glycolysis-PPP-TCA pathway discussed in the Results which includes the pentose phosphate pathway (PPP). Reducing the number of reactions limits the possible regulation schemes. When utilizing the same initial metabolite concentrations as the larger pathway when appropriate, i.e. high NAD/NADH (31.3), the regulation schemes for various methods show closer agreement. Both HEX1 and GAPD are regulated by every method as in the glycolysis-PPP-TCA pathway. The local MCA method, however, regulates PFK, PGK, and PDH, while the RL method additionally regulates PGI. Both methods regulate more reactions than the unrestriced MCA method and therefore result in a lower energy dissipation rate. Table S5 lists the complete reaction activity, flux and free energy for each respective prediction method.

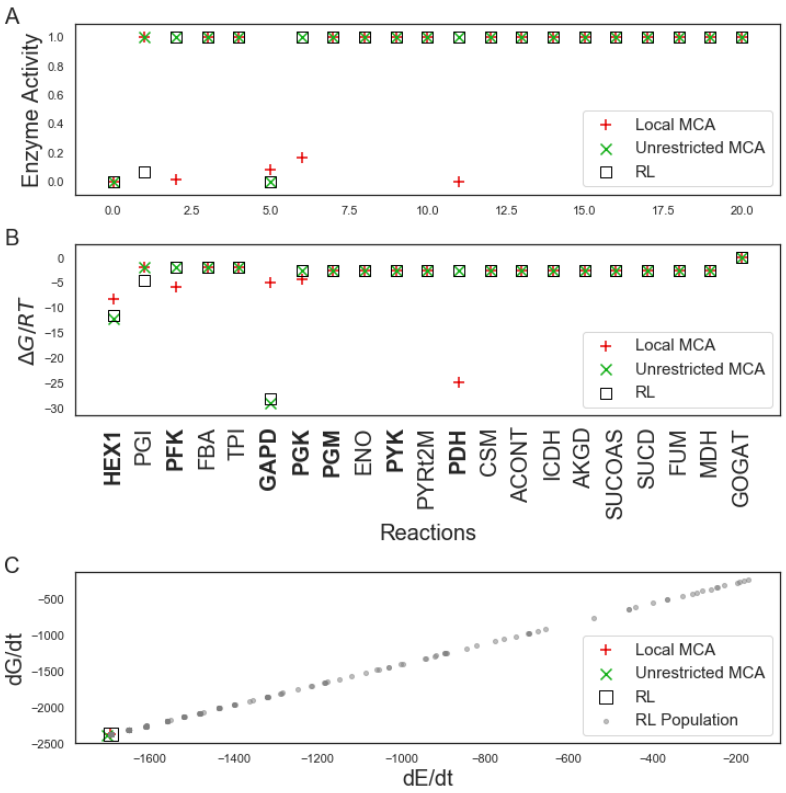


**Figure S3.** Glycolysis-TCA cycle predictions with high NAD/NADH initial conditions. Predicted enzyme activities (A) and free energy (B) at terminal states are calculated using concentration control theory, shown as red 'plus's and green 'X's, respectively. Results are compared to those found using a RL approach (black square). Grey dots (C) represent the population of terminal states found while training the RL agent.

### Tables

| NAD/NADH // NADP/NADPH Ratio | | High//Low | | | High//High | | | High//High no PFK | | |
|---|---|---|---|---|---|---|---|---|---|---|
| Method | | MCA Local | MCA | RL | MCA Local | MCA | RL | MCA Local | MCA | RL |
| Glycolysis | HEX1 | 6.34E+00 | 6.25E+00 | 6.28E+00 | 6.27E+00 | 8.23E+00 | 8.22E+00 | 5.76E-03 | 1.22E+01 | 1.22E+01 |
| | PGI | 6.34E+00 | 7.05E+00 | 7.21E+00 | 6.19E+00 | -2.80E+0 | -2.40E+00 | -1.15E-02 | -2.44E+01 | -2.44E+01 |
| | PFK | 6.34E+00 | 6.52E+00 | 6.59E+00 | 6.24E+00 | 4.56E+00 | 4.68E+00 | 0.00E+00 | 0.00E+00 | 0.00E+00 |
| | FBA | 6.34E+00 | 6.52E+00 | 6.59E+00 | 6.24E+00 | 4.56E+00 | 4.68E+00 | 1.05E-14 | -3.80E-15 | -3.80E-15 |
| | TPI | 6.34E+00 | 6.52E+00 | 6.59E+00 | 6.24E+00 | 4.56E+00 | 4.68E+00 | -7.80E-16 | 6.22E-15 | -7.80E-16 |
| | GAPD | 1.27E+01 | 1.28E+01 | 1.29E+01 | 1.25E+01 | 1.28E+01 | 1.29E+01 | 5.76E-03 | 1.22E+01 | 1.22E+01 |
| | PGK | 1.27E+01 | 1.28E+01 | 1.29E+01 | 1.25E+01 | 1.28E+01 | 1.29E+01 | 5.76E-03 | 1.22E+01 | 1.22E+01 |
| | PGM | 1.27E+01 | 1.28E+01 | 1.29E+01 | 1.25E+01 | 1.28E+01 | 1.29E+01 | 5.76E-03 | 1.22E+01 | 1.22E+01 |
| | ENO | 1.27E+01 | 1.28E+01 | 1.29E+01 | 1.25E+01 | 1.28E+01 | 1.29E+01 | 5.76E-03 | 1.22E+01 | 1.22E+01 |
| | PYK | 1.27E+01 | 1.28E+01 | 1.29E+01 | 1.25E+01 | 1.28E+01 | 1.29E+01 | 5.76E-03 | 1.22E+01 | 1.22E+01 |
| | PYRt2m | 1.27E+01 | 1.28E+01 | 1.29E+01 | 1.25E+01 | 1.28E+01 | 1.29E+01 | 5.76E-03 | 1.22E+01 | 1.22E+01 |
| | PDH | 1.27E+01 | 1.28E+01 | 1.29E+01 | 1.25E+01 | 1.28E+01 | 1.29E+01 | 5.76E-03 | 1.22E+01 | 1.22E+01 |
| PPP | G6PDH | 2.56E-14 | -7.98E-01 | -9.30E-01 | 8.09E-02 | 1.10E+01 | 1.06E+01 | 1.73E-02 | 3.66E+01 | 3.66E+01 |
| | PGL | 5.27E-33 | -7.98E-01 | -9.30E-01 | 8.09E-02 | 1.10E+01 | 1.06E+01 | 1.73E-02 | 3.66E+01 | 3.66E+01 |
| | GND | 3.18E-14 | -7.98E-01 | -9.30E-01 | 8.09E-02 | 1.10E+01 | 1.06E+01 | 1.73E-02 | 3.66E+01 | 3.66E+01 |
| | RPI | 8.10E-15 | -2.66E-01 | -3.10E-01 | 2.70E-02 | 3.68E+00 | 3.54E+00 | 5.76E-03 | 1.22E+01 | 1.22E+01 |
| | RPE | 1.42E-14 | -5.32E-01 | -6.20E-01 | 5.39E-02 | 7.36E+00 | 7.08E+00 | 1.15E-02 | 2.44E+01 | 2.44E+01 |
| | TKT1 | 4.88E-15 | -2.66E-01 | -3.10E-01 | 2.70E-02 | 3.68E+00 | 3.54E+00 | 5.76E-03 | 1.22E+01 | 1.22E+01 |
| | TALA | 9.77E-15 | -2.66E-01 | -3.10E-01 | 2.70E-02 | 3.68E+00 | 3.54E+00 | 5.76E-03 | 1.22E+01 | 1.22E+01 |
| | TKT2 | 1.07E-14 | -2.66E-01 | -3.10E-01 | 2.70E-02 | 3.68E+00 | 3.54E+00 | 5.76E-03 | 1.22E+01 | 1.22E+01 |
| TCA | CSM | 1.27E+01 | 1.28E+01 | 1.29E+01 | 1.25E+01 | 1.28E+01 | 1.29E+01 | 5.76E-03 | 1.22E+01 | 1.22E+01 |
| | ACONT | 1.27E+01 | 1.28E+01 | 1.29E+01 | 1.25E+01 | 1.28E+01 | 1.29E+01 | 5.76E-03 | 1.22E+01 | 1.22E+01 |
| | ICDH | 1.27E+01 | 1.28E+01 | 1.29E+01 | 1.25E+01 | 1.28E+01 | 1.29E+01 | 5.76E-03 | 1.22E+01 | 1.22E+01 |
| | AKGD | 1.27E+01 | 1.28E+01 | 1.29E+01 | 1.25E+01 | 1.28E+01 | 1.29E+01 | 5.76E-03 | 1.22E+01 | 1.22E+01 |
| | SUCOAS | 1.27E+01 | 1.28E+01 | 1.29E+01 | 1.25E+01 | 1.28E+01 | 1.29E+01 | 5.76E-03 | 1.22E+01 | 1.22E+01 |
| | SUCD | 1.27E+01 | 1.28E+01 | 1.29E+01 | 1.25E+01 | 1.28E+01 | 1.29E+01 | 5.76E-03 | 1.22E+01 | 1.22E+01 |
| | FUM | 1.27E+01 | 1.28E+01 | 1.29E+01 | 1.25E+01 | 1.28E+01 | 1.29E+01 | 5.76E-03 | 1.22E+01 | 1.22E+01 |
| | MDH | 1.27E+01 | 1.28E+01 | 1.29E+01 | 1.25E+01 | 1.28E+01 | 1.29E+01 | 5.76E-03 | 1.22E+01 | 1.22E+01 |
| | GOGAT | 6.99E-15 | 6.99E-15 | -1.10E-16 | -1.10E-16 | -1.10E-16 | -1.10E-16 | -1.10E-16 | -1.10E-16 | -1.10E-16 |

**Table S1.** Reaction fluxes at predicted enzyme activities from MCA-local, MCA, and RL methods for the glycolysis-PPP-TCA pathway under different boundary conditions.

| NAD/NADH // NADP/NADPH Ratio | | High//Low | | | High//High | | | High//High no PFK | | |
|---|---|---|---|---|---|---|---|---|---|---|
| Method | | MCA Local | MCA | RL | MCA Local | MCA | RL | MCA Local | MCA | RL |
| Glycolysis | HEX1 | -8.32E+00 | -1.63E+01 | -1.79E+01 | -8.31E+00 | -1.82E+01 | -2.00E+01 | -8.23E+00 | -2.28E+01 | -2.28E+01 |
| | PGI | -1.87E+00 | -1.97E+00 | -1.99E+00 | -1.85E+00 | 1.14E+00 | 1.01E+00 | 5.76E-03 | 3.19E+00 | 3.19E+00 |
| | PFK | -5.86E+00 | -1.90E+00 | -1.91E+00 | -5.85E+00 | -1.56E+00 | -1.59E+00 | -1.41E+02 | -5.49E+01 | -5.49E+01 |
| | FBA | -1.87E+00 | -1.90E+00 | -1.91E+00 | -1.86E+00 | -1.56E+00 | -1.59E+00 | -5.30E-15 | 1.78E-15 | 1.78E-15 |
| | TPI | -1.87E+00 | -1.90E+00 | -1.91E+00 | -1.86E+00 | -1.56E+00 | -1.59E+00 | 3.33E-16 | -3.10E-15 | 3.33E-16 |
| | GAPD | -4.99E+00 | -2.71E+01 | -1.04E+01 | -5.20E+00 | -2.82E+01 | -1.33E+01 | -2.88E-03 | -2.51E+00 | -2.51E+00 |
| | PGK | -4.32E+00 | -2.55E+00 | -9.92E+00 | -4.31E+00 | -2.55E+00 | -1.28E+01 | -2.88E-03 | -2.51E+00 | -2.51E+00 |
| | PGM | -2.55E+00 | -2.55E+00 | -2.56E+00 | -2.53E+00 | -2.55E+00 | -2.56E+00 | -2.88E-03 | -2.51E+00 | -2.51E+00 |
| | ENO | -2.55E+00 | -2.55E+00 | -2.56E+00 | -2.53E+00 | -2.55E+00 | -2.56E+00 | -2.88E-03 | -2.51E+00 | -2.51E+00 |
| | PYK | -2.55E+00 | -2.55E+00 | -5.90E+00 | -2.53E+00 | -2.55E+00 | -2.56E+00 | -2.88E-03 | -2.51E+00 | -2.51E+00 |
| | PYRt2m | -2.55E+00 | -2.55E+00 | -7.02E+00 | -2.53E+00 | -2.55E+00 | -6.13E+00 | -2.88E-03 | -2.51E+00 | -2.51E+00 |
| | PDH | -2.51E+01 | -2.55E+00 | -3.23E+00 | -2.51E+01 | -2.55E+00 | -2.56E+00 | -2.88E-03 | -2.51E+00 | -2.51E+00 |
| PPP | G6PDH | -1.30E-14 | 3.89E-01 | 4.50E-01 | -7.08E+00 | -2.41E+00 | -2.37E+00 | -7.10E+00 | -3.60E+00 | -3.60E+00 |
| | PGL | -3.84E-01 | 3.89E-01 | 4.50E-01 | -7.53E+00 | -2.41E+00 | -2.37E+00 | -4.20E+00 | -3.60E+00 | -3.60E+00 |
| | GND | -1.60E-14 | 3.89E-01 | 4.50E-01 | -4.04E-02 | -2.41E+00 | -2.37E+00 | -8.64E-03 | -3.60E+00 | -3.60E+00 |
| | RPI | -4.00E-15 | 1.33E-01 | 1.54E-01 | -1.35E-02 | -1.37E+00 | -1.34E+00 | -2.88E-03 | -2.51E+00 | -2.51E+00 |
| | RPE | -7.10E-15 | 2.63E-01 | 3.05E-01 | -2.70E-02 | -2.01E+00 | -1.98E+00 | -5.76E-03 | -3.19E+00 | -3.19E+00 |
| | TKT1 | -2.40E-15 | 1.33E-01 | 1.54E-01 | -1.98E+00 | -1.37E+00 | -1.34E+00 | -7.34E+01 | -2.51E+00 | -2.51E+00 |
| | TALA | -4.90E-15 | 1.33E-01 | 1.54E-01 | -1.35E-02 | -1.37E+00 | -1.34E+00 | -2.88E-03 | -2.51E+00 | -2.51E+00 |
| | TKT2 | -5.30E-15 | 1.33E-01 | 1.54E-01 | -1.35E-02 | -1.37E+00 | -1.34E+00 | -2.88E-03 | -2.51E+00 | -2.51E+00 |
| TCA | CSM | -2.55E+00 | -2.55E+00 | -2.56E+00 | -2.53E+00 | -2.55E+00 | -2.56E+00 | -2.88E-03 | -2.51E+00 | -2.51E+00 |
| | ACONT | -2.55E+00 | -2.55E+00 | -2.56E+00 | -2.53E+00 | -2.55E+00 | -2.56E+00 | -2.88E-03 | -2.51E+00 | -2.51E+00 |
| | ICDH | -2.55E+00 | -2.55E+00 | -2.56E+00 | -2.53E+00 | -2.55E+00 | -2.56E+00 | -2.88E-03 | -2.51E+00 | -2.51E+00 |
| | AKGD | -2.55E+00 | -2.55E+00 | -2.56E+00 | -2.53E+00 | -2.55E+00 | -2.56E+00 | -2.88E-03 | -2.51E+00 | -2.51E+00 |
| | SUCOAS | -2.55E+00 | -2.55E+00 | -2.56E+00 | -2.53E+00 | -2.55E+00 | -2.56E+00 | -2.88E-03 | -2.51E+00 | -2.51E+00 |
| | SUCD | -2.55E+00 | -2.55E+00 | -2.56E+00 | -2.53E+00 | -2.55E+00 | -2.56E+00 | -2.88E-03 | -2.51E+00 | -2.51E+00 |
| | FUM | -2.55E+00 | -2.55E+00 | -2.56E+00 | -2.53E+00 | -2.55E+00 | -2.56E+00 | -2.88E-03 | -2.51E+00 | -2.51E+00 |
| | MDH | -2.55E+00 | -2.55E+00 | -2.56E+00 | -2.53E+00 | -2.55E+00 | -2.56E+00 | -2.88E-03 | -2.51E+00 | -2.51E+00 |
| | GOGAT | -3.60E-15 | -3.60E-15 | 1.11E-16 | 1.11E-16 | 1.11E-16 | 1.11E-16 | 1.11E-16 | 1.11E-16 | 1.11E-16 |

**Table S2.** Reaction free energy at predicted enzyme activities from MCA-local, MCA, and RL methods for the glycolysis-PPP-TCA pathway under different boundary conditions.

| NAD/NADH // NADP/NADPH Ratio | | High//Low | | | High//High | | | High//High no PFK | | |
|---|---|---|---|---|---|---|---|---|---|---|
| Method | | MCA Local | MCA | RL | MCA Local | MCA | RL | MCA Local | MCA | RL |
| Glycolysis | HEX1 | 1.55E-03 | 5.02E-07 | 1.05E-07 | 1.55E-03 | 1.55E-03 | 1.77E-08 | 1.23E-06 | 1.52E-09 | 1.52E-09 |
| | PFK | 1.80E-02 | 1.00E+00 | 1.00E+00 | 1.80E-02 | 1.00E+00 | 1.00E+00 | 0.00E+00 | 0.00E+00 | 0.00E+00 |
| | GAPD | 8.59E-02 | 2.19E-11 | 4.06E-04 | 6.87E-02 | 7.17E-12 | 2.23E-05 | 1.00E+00 | 1.00E+00 | 1.00E+00 |
| | PGK | 1.68E-01 | 1.00E+00 | 6.34E-04 | 1.68E-01 | 1.00E+00 | 3.48E-05 | 1.00E+00 | 1.00E+00 | 1.00E+00 |
| | PYK | 1.00E+00 | 1.00E+00 | 3.52E-02 | 1.00E+00 | 1.00E+00 | 1.00E+00 | 1.00E+00 | 1.00E+00 | 1.00E+00 |
| | PYRt2m | 1.00E+00 | 1.00E+00 | 1.15E-02 | 1.00E+00 | 1.00E+00 | 2.81E-02 | 1.00E+00 | 1.00E+00 | 1.00E+00 |
| | PDH | 1.63E-10 | 1.00E+00 | 5.12E-01 | 1.63E-10 | 1.00E+00 | 1.00E+00 | 1.00E+00 | 1.00E+00 | 1.00E+00 |
| PPP | G6PDH | 1.00E+00 | 1.00E+00 | 1.00E+00 | 6.81E-05 | 1.00E+00 | 1.00E+00 | 1.43E-05 | 1.00E+00 | 1.00E+00 |
| | PGL | 5.36E-33 | 1.00E+00 | 1.00E+00 | 4.36E-05 | 1.00E+00 | 1.00E+00 | 2.60E-04 | 1.00E+00 | 1.00E+00 |
| | TKT1 | 1.00E+00 | 1.00E+00 | 1.00E+00 | 3.78E-03 | 1.00E+00 | 1.00E+00 | 7.72E-35 | 1.00E+00 | 1.00E+00 |

**Table S3.** Predicted enzyme activities from MCA-local, MCA, and RL methods for the glycolysis-PPP-TCA pathway under different boundary conditions. Unlisted reactions are unregulated.

| | | Activity | | | Flux | | | Energy | | |
|---|---|---|---|---|---|---|---|---|---|---|
| Method | | MCA Local | MCA | RL | MCA Local | MCA | RL | MCA Local | MCA | RL |
| Gluconeogenesis | G6Pase | 1.00E+00 | 1.00E+00 | 1.00E+00 | 6.33E-01 | 6.33E-01 | 6.33E-01 | -3.11E-01 | -3.11E-01 | -3.11E-01 |
| | PGI | 1.00E+00 | 1.00E+00 | 1.00E+00 | 6.33E-01 | 6.33E-01 | 6.33E-01 | -3.11E-01 | -3.11E-01 | -3.11E-01 |
| | FBP | 1.00E+00 | 1.00E+00 | 1.00E+00 | 6.33E-01 | 6.33E-01 | 6.33E-01 | -3.11E-01 | -3.11E-01 | -3.11E-01 |
| | FBA | 1.00E+00 | 1.00E+00 | 1.00E+00 | 6.33E-01 | 6.33E-01 | 6.33E-01 | -3.11E-01 | -3.11E-01 | -3.11E-01 |
| | GAPD | 1.00E+00 | 1.00E+00 | 1.00E+00 | 1.27E+00 | 1.27E+00 | 1.27E+00 | -5.97E-01 | -5.97E-01 | -5.97E-01 |
| | PGK | 1.00E+00 | 1.00E+00 | 1.00E+00 | 1.27E+00 | 1.27E+00 | 1.27E+00 | -5.97E-01 | -5.97E-01 | -5.97E-01 |
| | PGM | 1.00E+00 | 1.00E+00 | 1.00E+00 | 1.27E+00 | 1.27E+00 | 1.27E+00 | -5.97E-01 | -5.97E-01 | -5.97E-01 |
| | ENO | 1.00E+00 | 1.00E+00 | 1.00E+00 | 1.27E+00 | 1.27E+00 | 1.27E+00 | -5.97E-01 | -5.97E-01 | -5.97E-01 |
| | PEPCK | 1.00E+00 | 1.00E+00 | 1.00E+00 | 1.27E+00 | 1.27E+00 | 1.27E+00 | -5.97E-01 | -5.97E-01 | -5.97E-01 |
| | PC | 9.90E-04 | 9.90E-04 | 9.90E-04 | 1.27E+00 | 1.27E+00 | 1.26E+00 | -7.15E+00 | -7.15E+00 | -7.15E+00 |

**Table S4.** Predicted enzyme activities, fluxes, and free energy for gluconeogenesis pathway from MCA-local, MCA, and RL methods.

| | | Activity | | | Flux | | | Energy | | |
|---|---|---|---|---|---|---|---|---|---|---|
| Method | | MCA Local | MCA | RL | MCA Local | MCA | RL | MCA Local | MCA | RL |
| Glycolysis | HEX1 | 1.55E-03 | 3.48E-05 | 6.81E-05 | 6.42E+00 | 6.45E+00 | 6.41E+00 | -8.33E+00 | -1.21E+01 | -1.15E+01 |
| | PGI | 1.00E+00 | 1.00E+00 | 6.87E-02 | 6.42E+00 | 6.45E+00 | 6.41E+00 | -1.88E+00 | -1.89E+00 | -4.54E+00 |
| | PFK | 1.80E-02 | 1.00E+00 | 1.00E+00 | 6.42E+00 | 6.45E+00 | 6.41E+00 | -5.88E+00 | -1.89E+00 | -1.88E+00 |
| | FBA | 1.00E+00 | 1.00E+00 | 1.00E+00 | 6.42E+00 | 6.45E+00 | 6.41E+00 | -1.88E+00 | -1.89E+00 | -1.88E+00 |
| | TPI | 1.00E+00 | 1.00E+00 | 1.00E+00 | 6.42E+00 | 6.45E+00 | 6.41E+00 | -1.88E+00 | -1.89E+00 | -1.88E+00 |
| | GAPD | 8.59E-02 | 2.94E-12 | 7.17E-12 | 1.28E+01 | 1.29E+01 | 1.28E+01 | -5.01E+00 | -2.91E+01 | -2.82E+01 |
| | PGK | 1.68E-01 | 1.00E+00 | 1.00E+00 | 1.28E+01 | 1.29E+01 | 1.28E+01 | -4.34E+00 | -2.56E+00 | -2.56E+00 |
| | PGM | 1.00E+00 | 1.00E+00 | 1.00E+00 | 1.28E+01 | 1.29E+01 | 1.28E+01 | -2.56E+00 | -2.56E+00 | -2.56E+00 |
| | ENO | 1.00E+00 | 1.00E+00 | 1.00E+00 | 1.28E+01 | 1.29E+01 | 1.28E+01 | -2.56E+00 | -2.56E+00 | -2.56E+00 |
| | PYK | 1.00E+00 | 1.00E+00 | 1.00E+00 | 1.28E+01 | 1.29E+01 | 1.28E+01 | -2.56E+00 | -2.56E+00 | -2.56E+00 |
| | PYRt2m | 1.00E+00 | 1.00E+00 | 1.00E+00 | 1.28E+01 | 1.29E+01 | 1.28E+01 | -2.56E+00 | -2.56E+00 | -2.56E+00 |
| | PDH | 2.04E-10 | 1.00E+00 | 1.00E+00 | 1.28E+01 | 1.29E+01 | 1.28E+01 | -2.49E+01 | -2.56E+00 | -2.56E+00 |
| TCA | CSM | 1.00E+00 | 1.00E+00 | 1.00E+00 | 1.28E+01 | 1.29E+01 | 1.28E+01 | -2.56E+00 | -2.56E+00 | -2.56E+00 |
| | ACONT | 1.00E+00 | 1.00E+00 | 1.00E+00 | 1.28E+01 | 1.29E+01 | 1.28E+01 | -2.56E+00 | -2.56E+00 | -2.56E+00 |
| | ICDH | 1.00E+00 | 1.00E+00 | 1.00E+00 | 1.28E+01 | 1.29E+01 | 1.28E+01 | -2.56E+00 | -2.56E+00 | -2.56E+00 |
| | AKGD | 1.00E+00 | 1.00E+00 | 1.00E+00 | 1.28E+01 | 1.29E+01 | 1.28E+01 | -2.56E+00 | -2.56E+00 | -2.56E+00 |
| | SUCOAS | 1.00E+00 | 1.00E+00 | 1.00E+00 | 1.28E+01 | 1.29E+01 | 1.28E+01 | -2.56E+00 | -2.56E+00 | -2.56E+00 |
| | SUCD | 1.00E+00 | 1.00E+00 | 1.00E+00 | 1.28E+01 | 1.29E+01 | 1.28E+01 | -2.56E+00 | -2.56E+00 | -2.56E+00 |
| | FUM | 1.00E+00 | 1.00E+00 | 1.00E+00 | 1.28E+01 | 1.29E+01 | 1.28E+01 | -2.56E+00 | -2.56E+00 | -2.56E+00 |
| | MDH | 1.00E+00 | 1.00E+00 | 1.00E+00 | 1.28E+01 | 1.29E+01 | 1.28E+01 | -2.56E+00 | -2.56E+00 | -2.56E+00 |
| | GOGAT | 1.00E+00 | 1.00E+00 | 1.00E+00 | -1.10E-16 | -1.10E-16 | -1.10E-16 | 1.11E-16 | 1.11E-16 | 1.11E-16 |

**Table S5.** Predicted enzyme activities, fluxes, and free energy for glycolysis-TCA pathway from MCA-local, MCA, and RL methods.
